# NECing goes: flexibility of the herpesvirus nuclear egress complex

**DOI:** 10.1101/2023.07.07.547920

**Authors:** Vojtěch Pražák, Yuliia Mironova, Daven Vasishtan, Christoph Hagen, Ulrike Laugks, Yannick Jensen, Saskia Sanders, John M. Heumann, Jens B. Bosse, Barbara Klupp, Thomas C. Mettenleiter, Michael Grange, Kay Grünewald

## Abstract

The nuclear egress complex (NEC) allows herpesvirus capsids to escape from the nucleus without breaking the nuclear envelope barrier. It assembles into a lattice on the inner nuclear membrane enveloping newly assembled nucleocapsids, which bud into the perinuclear space. The primary virion envelope subsequently fuses with the outer nuclear membrane, releasing the capsid into the cytosol. Here we interrogated the NEC in the context of intact cells infected with pseudorabies or herpes simplex virus using focused-ion beam milling and electron cryo- tomography. We determined the structure of NEC in different conformations and show that it consists of a flexible hexameric lattice that generates curvature through a combination of ordered and disordered domains. After interrogating the intermediate stages of capsid formation, we show that capsid vertex binding may initiate envelopment but does not directly induce curvature formation. These data and many examples of the intermediate stages of nuclear egress paint a detailed holistic view of a versatile transport system.

## Introduction

Mammalian cells contain a number of membrane-bound and membrane-less organelles that allow the spatiotemporal organisation of macromolecules according to their function. This compartmentalisation increases the efficiency of processes confined within. As a consequence, cells have evolved a number of different transport mechanisms to move material to and from the cellular space in which they are required. The ability for viruses to manipulate and subvert these cellular machineries is essential for the transport of viral gene products within cells and the eventual production of viral progeny. Herpesviruses are no exception. Upon fusing with the cellular plasma membrane, herpesvirus capsids enter the cytosol and are then trafficked to the nuclear membrane^1^. Interaction with the nuclear pore complex (NPC) results in the injection of the viral DNA into the nucleoplasm, enabling its replication and subsequent production of viral mRNA required for production of viral proteins^2^.

Nucleocapsids are assembled within the nucleoplasm, and once produced exceed the ∼40 nm size exclusion limit of the NPC^2^ (nucleocapsids have a diameter of ∼120 nm). Consequently, nucleocapsids are transported across the inner and outer nuclear membranes (INM and ONM) via a specialised mechanism involving membrane envelopment/de-envelopment termed herpesvirus nuclear egress^3, 4^. Firstly, newly assembled nucleocapsids associate with, and are enveloped at, the INM to produce primary-enveloped (perinuclear) particles residing in the perinuclear space (the lumen between the INM and ONM). Secondly, the primary virion envelope fuses with ONM to release the nucleocapsids into the cytosol for subsequent stages of assembly and egress^3^. A similar process was later described in Drosophila cells, where large protein aggregates from abortive processes within the nucleus are transported across the INM and ONM for degradation by the cytosolic autophagic machinery^5^.

A complex fibrous protein network called the nuclear lamina exists underneath the INM which is composed of A-/C- and B-type lamins as well as other membrane-bound proteins. This network must be disrupted locally in order to clear space for egress to occur^6^. Herpesviruses encode two proteins, designated pUL34 and pUL31 (in the case of the alphaherpesviruses such as pseudorabies virus, PrV, and herpes simplex virus-1, HSV1), that are the two main mediators of herpesvirus nuclear egress. pUL31 is a globular phosphoprotein that together with pUL34 (a single pass transmembrane protein) forms heterodimers at the INM, known as the nuclear egress complex (NEC)^7^. pUL31, pUL34 and the alphaherpesvirus-specific protein kinase pUS13 are together involved in a phosphorylation cascade including protein kinase A and protein kinase C^8, 9^ that leads to the local dissolution of the nuclear lamina. The heterodimers of pUL31/34 self-associate on the INM and subsequently cause invaginations upon association with a nucleocapsid. When expressed in eukaryotic cells in the absence of other viral proteins, the NEC is able to form hexameric lattices that induce specific curvature to the INM resulting in the formation of perinuclear vesicles^10–12^.

The NEC has been observed as an electron-dense coat between the INM and the nucleocapsid^10^. However, the precise mechanism of how nucleocapsids are recruited by the NEC remains ambiguous. A number of candidates, besides pUL31, have been identified that may mediate the physical interaction between nucleocapsid and NEC/INM. Chief among these candidates are pUL17 and pUL25^13–15^ even though pUL25 is not strictly required for egress^16^, which together with pUL36 form a structural component of the nucleocapsid termed the capsid-vertex specific component (CVSC)^17–19^. The CVSC rests on the five-fold vertex of the nucleocapsid of herpesviruses and has downstream functions in the recruitment of viral tegument once the nucleocapsid has crossed the nuclear membrane^20, 21^. Whether the CVSC specifically mediates the NEC-capsid interaction remains to be determined. Similarly, how these proteins function to regulate the type of capsid recruited into primary particles is an open question.

The NEC is able to form roughly spherical vesicles in the absence of nucleocapsid ^10^ suggesting that perhaps the nucleocapsid has a limited role in the formation of primary vesicles. The observation that the NEC assembles in a hexagonal geometry creates a problem when considering how the NEC may be able to form spherical structures, as this would require either the presence of pentons or a breakdown in the lattice. The pUL31/34 homologous heterodimeric X-ray crystallographic structures (and an NMR structure) have been solved previously for herpes simplex virus-1 (HSV1), pseudorabies virus (PrV), and a number of other members of Herpesviridae^22–28^ In the case of HSV1 and HCMV, the structures were solved in a P6 symmetry, allowing a molecular analysis of the interface between adjacent heterodimers in a (albeit flat) hexameric lattice. However, these data were insufficient to construct a mechanistic model of how curvature is induced. Isolated pUL25 can force NEC into a pentameric lattice in vitro but this interaction is yet to be confirmed in a more holistic system^29^. Recently, it has been shown that the NEC membrane proximal region can directly induce membrane curvature^30^ but it is unlikely to be the sole curvature inducing mechanism. Finally, in order to release the nucleocapsid into the cytosol, the primary vesicle membrane must fuse with the ONM. So far, no viral or cellular protein has been shown responsible this process, although it has been suggested in alphaherpesviruses to be enhanced by the protein pUS3^13, 31–33^, which functions as a central kinase in a number of different processes in the assembly and egress of herpesviruses^34^. Depletion of US3 from HSV1 and PrV viruses leads to an accumulation of primary particles in the perinuclear space, and a reduction in the amount of capsid released into the cytosol^31, 32^. The active machinery mediating fusion with the ONM is yet to be determined. It has been suggested that either gB or gH are required for nuclear egress of HSV (with gB being the only confirmed fusogen^35^. On the other hand, none of the entry glycoproteins are required for PrV egress^36^. As such, more work needs to be done to ascertain the true fusogen that mediates de-envelopment of primary particles in the perinuclear space.

Previous studies on herpesvirus nuclear egress and the interaction between the NEC and capsid have relied heavily on fluorescence microscopy, conventional transmission electron microscopy and biochemical analyses that are unable to highlight the molecular and structural details of this process to resolutions greater than ∼5 nm ^12, 16, 37, 38^. These techniques often require the fixation, and hence alteration, of cellular material, making the interpretation of specific subcellular structures problematic. Other attempts at determining the structure of the NEC in association with the herpesvirus nucleocapsid have required the isolation of material from cells in such a way as to alter their molecular arrangement, similarly making interpretation difficult^39, 40^. Recent developments in the field of electron cryo-microscopy (cryoEM), electron cryo-tomography (cryoET) and focused ion beam/electron scanning cryo-microscopy (cryoFIB-SEM)^41^ have allowed the assessment of ultrastructural and molecular details of biological material, from cell culture to tissues in a close to native state, thus facilitating interpretation of molecular details and allowing downstream structural analyses.

In the present study, we report the structure of the NEC and its interaction with nuclear capsids in primary enveloped (perinuclear) particles in the context of a PrV-ΔUS3, wild-type (WT) PrV, and HSV infection. In the PrV-ΔUS3 system, an accumulation of primary particles in the perinuclear space occurs, enabling sufficient volumes of data to be acquired for high resolution structural analyses. We image capsids egressing from the cell nucleus in a frozen hydrated state, and directly visualise capsids during different stages of primary envelopment/de-envelopment. We then perform structural analyses, primarily subvolume averaging, to investigate the structures of the NEC, newly formed capsids, and the interaction between them.

We explore how the initial interaction between newly formed capsids and the NEC leads to primary envelopment. We show that CVSC is unlikely to mediate this interaction as it is only fully assembled in cytosol. To produce perinuclear vesicles, the NEC assembles into a flexible hexameric array that induces budding of roughly spherical vesicles through multiple elongated patches separated by regions of disorder. A comparison between pseudoatomic models based on structures with different curvatures allowed us to directly show that the hexameric rings that form the basis of the NEC lattice are highly flexible, are capable of taking different, less symmetrical forms. Rather than changes within the hexamer, it is the interaction between neighbouring hexamers at the membrane proximal region of pUL34 that is likely the structural basis of curvature formation. Together, we present a holistic view of herpesvirus nuclear egress and a basis for integrating targeted perturbations of this transport system.

## Results

### CryoFIB-SEM and cryoET reveal a number of canonical and non-canonical structures during pseudorabies virus nuclear egress in-situ

We used cryoFIB-SEM thinning of porcine cells infected with PrV-ΔUS3 and subsequent imaging by cryoET to directly view the molecular details of herpesvirus assembly and nuclear egress at multiple stages (Fig. 1, S1). We noticed several structures relating to capsid assembly. Roughly spherical halos within the nucleoplasm that resembled the scaffold-ring within adjacent spherical pro-capsids and icosahedral B-capsids were visible (Fig. 1b, c, Movie 1)^42^.

**Figure 1.**
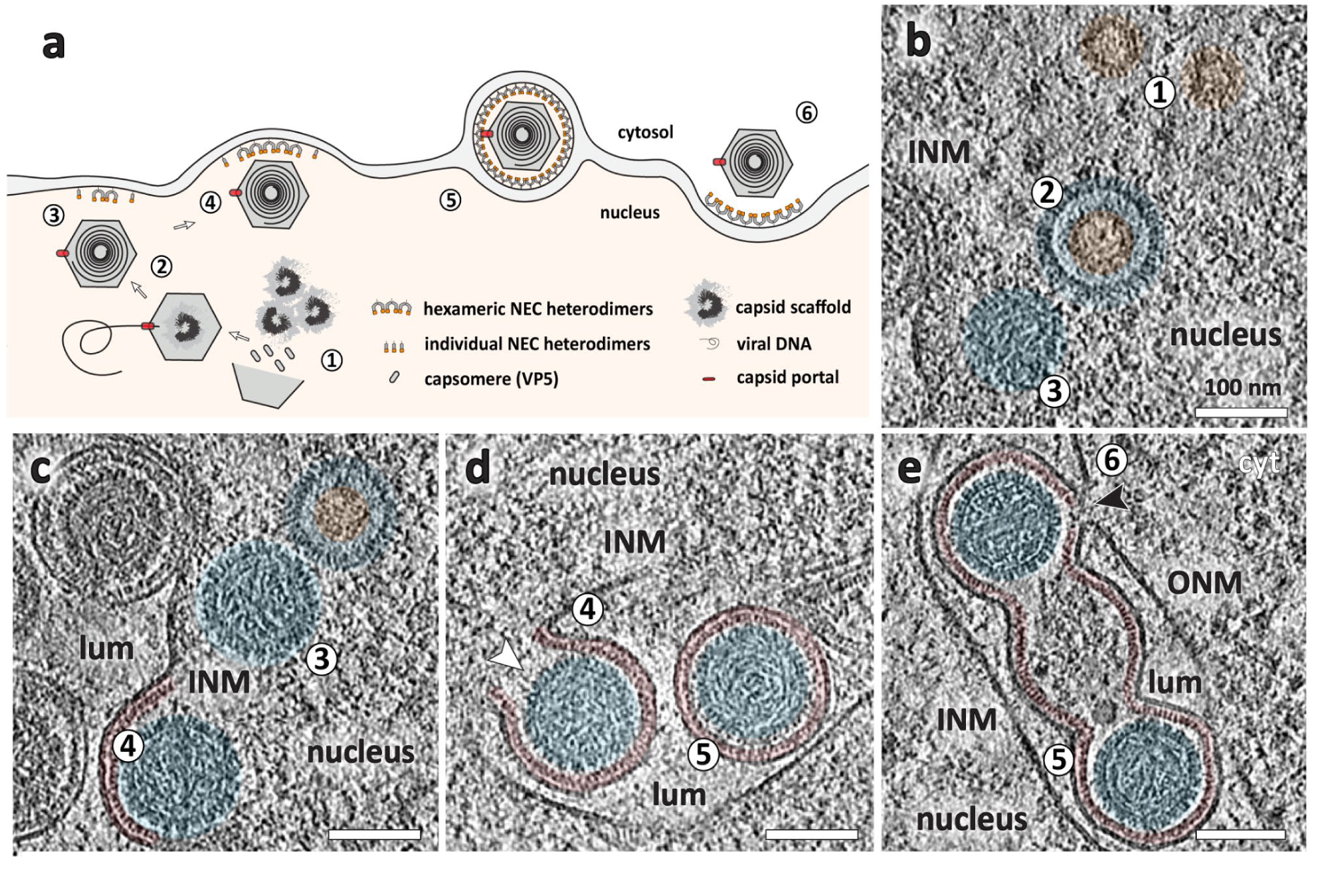
Visualising stages of PrV nuclear egress. **a** Schematic model of procapsid assembly and nuclear capsid egress via primary particle formation in the perinuclear space. Numbers indicate (selected) consecutive stages of nuclear egress. **b** CryoET slice illustrating stages 1-3 of nuclear egress, where 1 shows pro-capsid scaffold (orange), 2 shows a pro-/B-capsid in the nucleus (blue, with orange pro-capsid scaffold inside) and 3 showing an assembled C-capsid (blue). **c** CryoET slice of stage 3 and 4 leading up to interaction of nuclear C-capsid (blue) with nuclear egress complex associated with the inner nuclear membrane (red). **d** Budding of nucleocapsid into the perinuclear space (lumen between the INM and ONM) to form primary virions (stages 4 and 5). White arrow indicates membrane opening to the nucleus. **e** Primary virions in the lumen/perinuclear space traversing across the INM and ONM (stage 5 and 6). Black arrow indicates membrane of the primary vesicle open to the cytosol. Scale bars are 100 nm. ONM = outer nuclear membrane, INM = inner nuclear membrane, lum = lumen/perinuclear space.

The process of nucleocapsid envelopment into primary vesicles, and subsequent egress, was seen directly at several stages. At the INM, we observed a ∼10 nm thick NEC coat, in a manner consistent with previous studies^10^ (Fig. 1c).The NEC coat could be observed in association of nucleocapsids at the INM, with localised patches visibly causing inward curvature of the membrane (Fig. 1c, e). Primary particles could be observed at different stages of budding, from (presumably) initial nucleocapsid-NEC contact to almost fully formed vesicles still connected to the nucleoplasm through a narrow pore (Fig. 1d). Fully formed perinuclear vesicles, roughly spherical in shape, isolated or in clusters, constituted the majority of the NEC lattice occurrences in our data. The mean inner diameter of 163 isolated perinuclear vesicles was 119 +/- 10 nm, which is ∼20 nm larger than empty vesicles generated by expression of pUL31/34 alone^10^ (Fig. S2) Finally, there were primary vesicles partially opened to the cytosol, with fusion pores visible in the ONM, exposing the NEC coat to the cytosolic environment. Interestingly, the NEC lattice remained intact while the ONM was clearly open to the cytosol (Fig. 1e, S1, Movie 3).

Apart from singular, well-formed primary virions which had budded individually into the perinuclear lumen, we also observed enlarged, type-1 nucleoplasmic reticula-like^43^ (NR), structures containing several primary vesicles in small bundles, similar to previously described herniations^44, 45^ (Fig. 2b, c, Supp. Movie 4). We could also observe type-2 NR, in which invaginations of both INM and ONM allowed the close proximity of cytosol to the centre of nuclei^43^ (Fig. 2d, S1e). The primary vesicles in type 1-NR could sometimes be seen to form openings between several other primary particles and form interconnected compartments, all of which were supported by NEC, even in the absence of cargo (Fig. 1e). Indeed, we also observed elongated perinuclear vesicles/tubes that contained pro-capsid assembly structures (Fig. 2c, e, g). One very striking observation was the propensity for the NEC to also form extensive, elongated tubular structures, often devoid of large cargo but sometimes containing pro-capsid scaffolds or capsids, which propagated within the perinuclear lumen (Movie 2). Finally, there were several instances of NEC lattice interacting head to head and even forming flat, double-layer sheets (Fig. 2f, g, S3). These observations show that the NEC is capable of forming a variety of distinct structures depending on the molecular context and can transport capsids at different stages of assembly.

**Figure 2.**
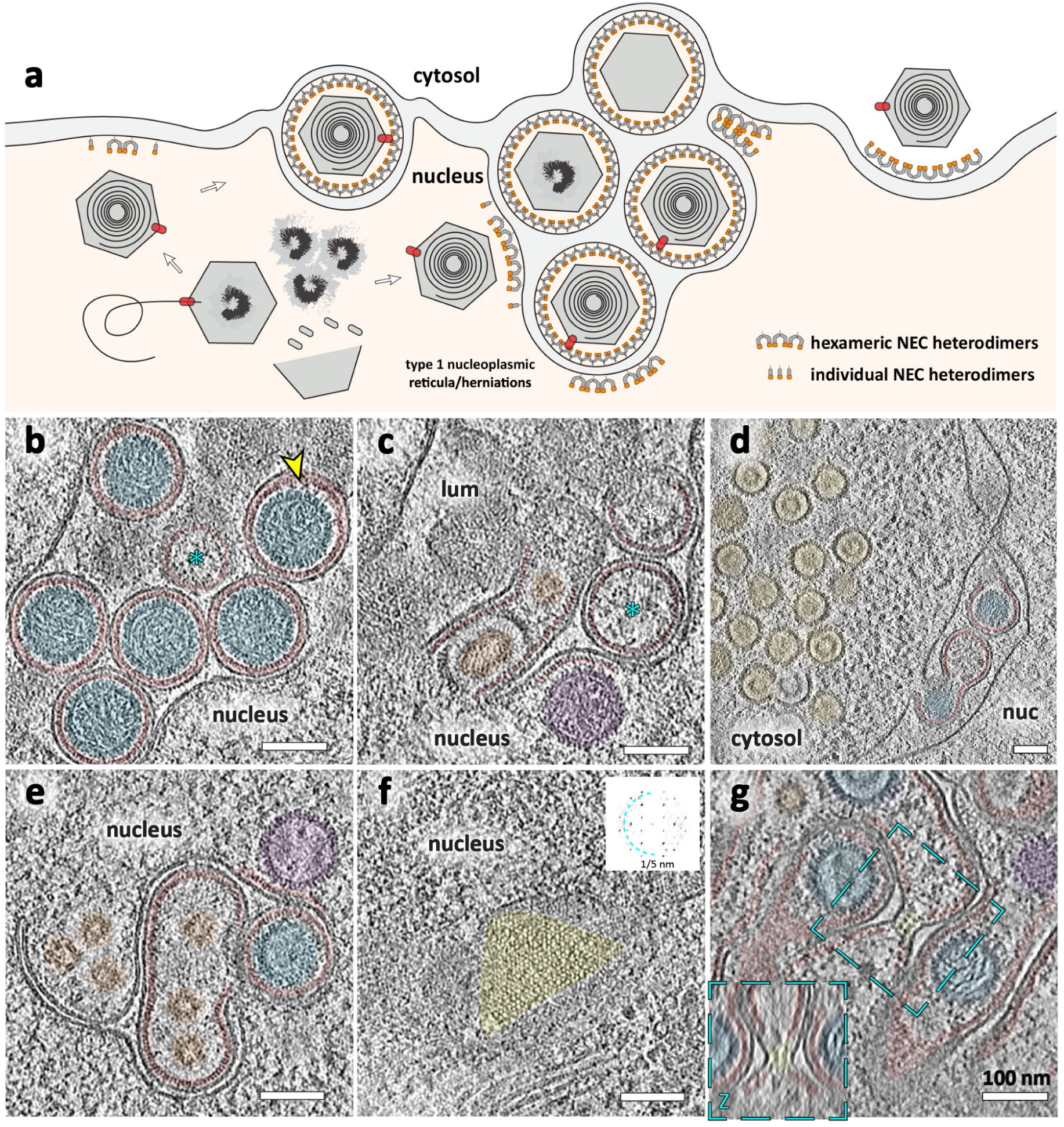
Visualisation of non-canonical stages/structures of PrV-ΔUS3 nuclear egress. **a** Schematic model of non-canonical nuclear egress. **b** CryoET slice of perinuclear, primary virions (blue) inside a large herniation within the lumen between INM and ONM. Red = nuclear egress complex. A cyan asterisk notes an empty perinuclear vesicle, (i.e. lacking assembled capsid components; it is not sectioned through the equator). Yellow arrow indicates an example of a vesicle with a gap between NEC and capsid. **c** A broken capsid entering NEC tubular herniation, with an additional pro-capsid scaffold (orange) and empty perinuclear vesicles (cyan asterisks). **d** A clump of partially assembled procapsids and B-capsids in type-2 nucleoplasmic reticulum (cytosol). **e** A perinuclear vesicle and an INM infolding containing several pro-capsid scaffolds (orange). **f** Top view of a putative head-head stacked double layer of pUL31/34 (yellow). Inset shows the power spectrum of a projection through the lattice layer. **g** Slice through several adjacent type-1 NR, with the lumen of one of these zippered by putative head-head interacting pUL31/34 (yellow). Inset shows an orthogonal slice through the centre of the highlighted area. Scales bars represent 100 nm.

**Figure 3.**
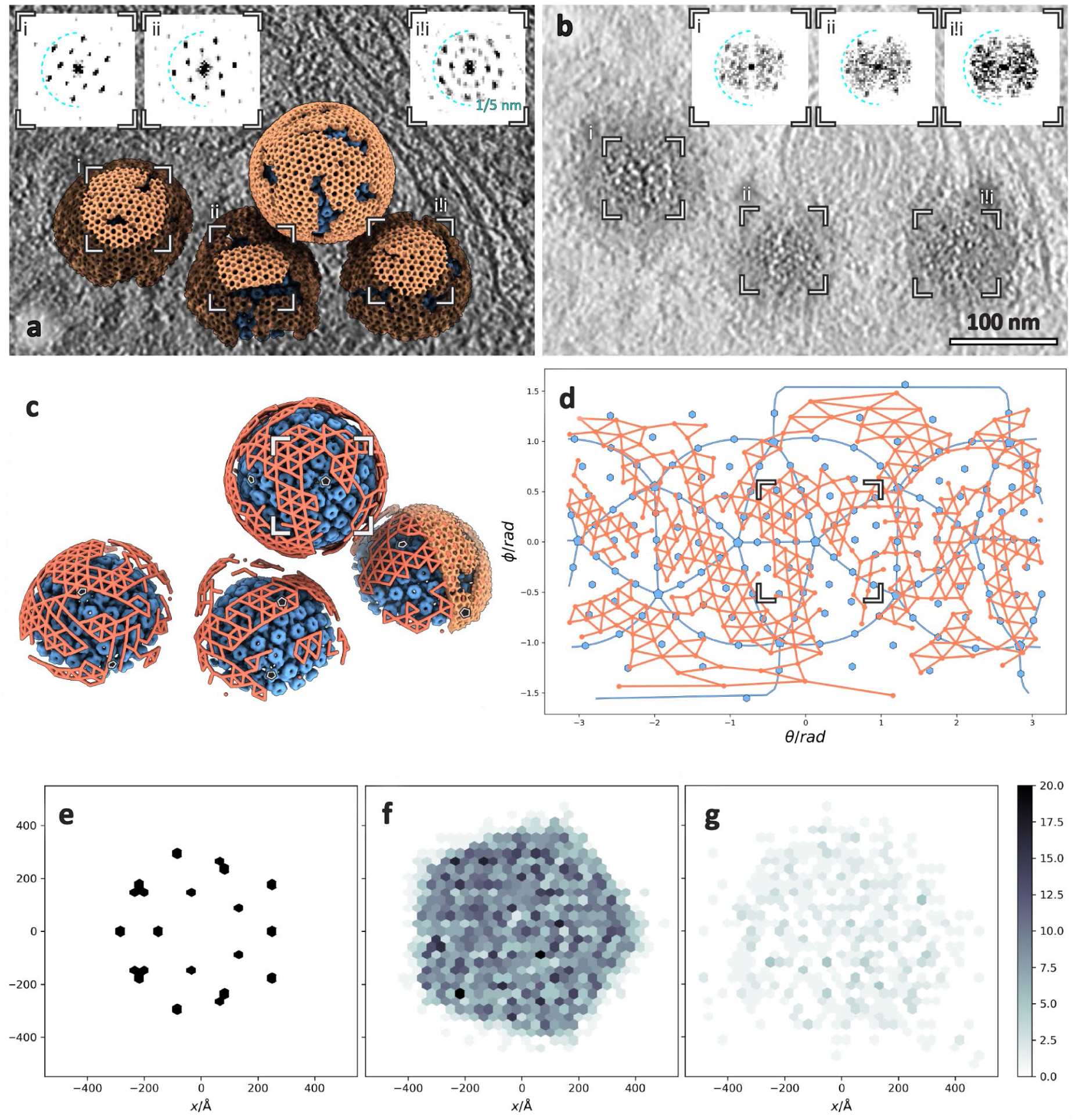
PrV NEC forms a locally ordered lattice. **a** 3D rendering of the NEC and nucleocapsids generated by backplotting subvolume averaging maps. Insets show contrast adjusted power spectra of sections through the highlighted areas intended for comparison to the raw data shown in b. **b** Section through raw tomogram showing tangential views of 3 perinuclear NEC vesicles. Insets show power spectra of the highlighted areas. iii is the result of twinning of two hexagonal symmetrical regions. **c** The same particles, but with NEC represented schematically as lines connecting hexamer centres. Some capsid vertices have been indicated with pentamers. **d** Polar coordinate representation of c, with origin at the nucleocapsid centre. Blue lines represent icosahedral edges. **e, f, g** Histogram of the relative distribution of hexons (e, used as a control), perinuclear NEC hexamers (f) and budding NEC hexamers (g) relative to pentons vertices. Scale bar represents the number of particles in each bin. As expected, hexon positions show a regular arrangement relative to the five-fold vertex, whereas the NEC positions appear to be randomly distributed.

### NEC-capsid contacts are pleomorphic

While we observed infrequent perinuclear vesicles without identifiable cargo, the majority of vesicles contained nucleocapsids. This suggests that there is a (more or less) specific NEC- nucleocapsid interaction that triggers perinuclear vesicle formation, if not also the initial assembly of pUL31/34 into a curvature-inducing lattice. Since both NEC and nucleocapsids are periodic structures, their specific interaction could manifest as an alignment between their respective geometries. Towards this end, we determined the structures of the NEC hexamer and capsid pentons and hexons in perinuclear vesicles using subvolume averaging (see methods). This provided us with their structures, but it also accurately and empirically located their positions, allowing us to analyse the correlation between them. First, to verify that these measurements accurately represent the lattice geometry, we placed NEC hexamers back into the original volume (backplotted) and compared the resulting model to the original data; the two were a good match both in real and reciprocal space (Fig. 3a, b). The hexameric NEC formed ordered domains across the surface of the vesicles, with patches of disorder evident in between the areas where the hexagonal symmetry persisted (Fig. 3c, d). A similar phenomenon that has been previously seen for the hexameric lattices produced by the Gag protein found on the inside of immature HIV-1 virions^46, 47^. As it is impossible to tile a sphere with hexagonal units, we hypothesise that the NEC forms spherical vesicles through locally curved sheets separated by areas of disorder. Consistent with our hypothesis, we observed elongated hexagonal patches often more than 2 radians long but only up to 1 radian in the orthogonal direction. Curvature in hexagonal arrays is allowed in one direction without deformation, while a simultaneous curvature in the orthogonal direction results in increasing amounts of disorder. Another way of approximating a spherical surface is using thin tubular sections (like a volleyball), but the analysis of local curvature within ordered regions suggest that they are curved in all directions (Fig S4). Whether the ordered patches are relatively stable after formation (akin to tectonic plates) or whether there is a dynamic exchange of subunits between areas of order remains to be determined.

Areas of NEC disorder do not preclude an alignment of the hexameric regions with the capsid icosahedral symmetry. On the contrary, they are required in the absence of NEC pentamers that would aid the formation of spherical vesicles. In the current study we could not discern any pentamers formed by the NEC. Visually comparing individual nucleocapsid and NEC lattices, we noticed no consistent pattern in the alignment of the two (Fig. 3c, d). Similarly, a global analysis of the distribution of NEC hexamers relative to capsid pentons showed no measurable correlation of the distance and orientation between these structures (Fig. 3e, f).

These data suggest that the NEC does not align itself with nucleocapsid symmetry in fully formed perinuclear vesicles. However, it could be the case that only a single vertex, for example, triggers NEC lattice formation, which then extends irrespective of the nature of the cargo. In this case, the lattice alignment of a single vertex would be disguised by the other 11 vertices. With this in mind, we focused our attention on nuclear capsids in proximity to the INM, which could be in the initial stages of envelopment (Fig. 4). In the majority of cases (19 out of 28) where nuclear capsids came within 20 nm of the INM, we could see patches of NEC already assembled in a hexameric array, as evidenced by strong six-fold peaks in their power spectra (Fig. 4d). These patches had a positive curvature, similar to that of fully formed perinuclear vesicles. Assessing these data globally (Fig. 3g) and on an individual case basis (Fig. S5, S6) again showed no correlation between the respective symmetries.

**Figure 4.**
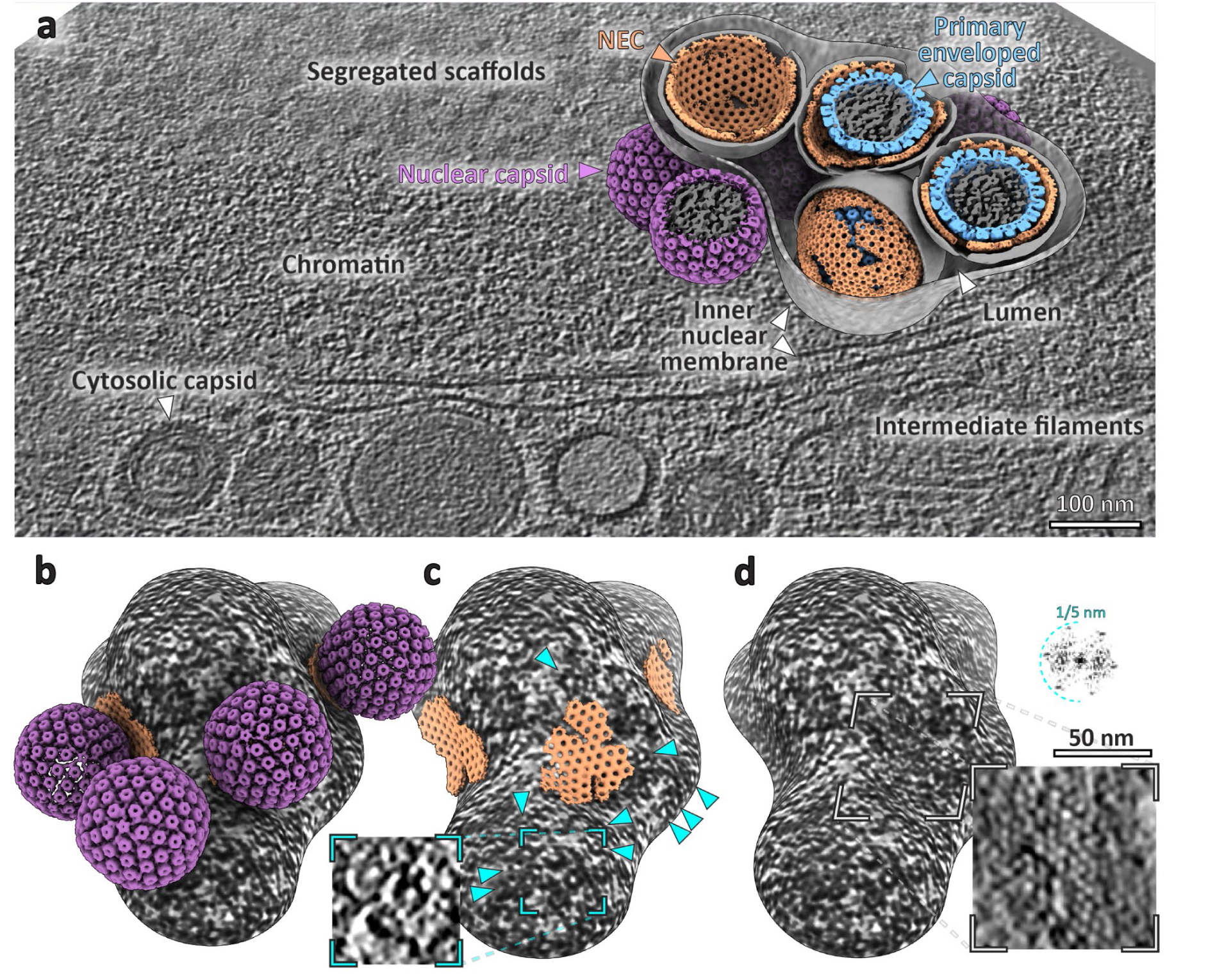
PrV nuclear egress complex in context. a. A slice through a tomogram and backplotted 3D representation of egress components. The membrane of one NEC vesicle has been cut away to expose its contents, similarly, one nucleocapsid volume has been omitted to show the inner NEC surface. **b, c, d** The same rendering, but flipped upside-down to show the detail of assembling NEC and capsid interaction. Cyan arrowheads in c indicate putative ring-like NEC located on negatively-curved membrane (inset shows a slice through the highlighted volume, same scale as d). Insets in d show a slice through the raw tomogram and the power spectrum of the highlighted area.

Multiple studies have suggested that the pUL25/pUL17 two proteins complex at the penton vertex, are needed to trigger egress (either directly or indirectly); however, conceptually there is no fundamental reason why the NEC should align itself with the nucleocapsid icosahedral symmetry and indeed, the main capsid protein (VP5) and pUL31 have complimentary electrostatics ^24^. To interrogate our dataset for hints of contacts, we classified our dataset of putative budding events based on the penton-NEC distance. Having obtained accurate positions and orientations of nucleocapsid pentons, we extracted and averaged the volumes of nearby NEC and vice versa (Fig. S7). The resulting densities can be interpreted similarly to the coordinate histogram representation (3e-g) with the important caveat being that the density does not depend on specifying reference coordinates relative to either lattice. Again, there was no evidence of pentons being aligned with any part of the NEC lattice. Weak bridging densities originating at penton vertices at ∼ 7 nm separation could be observed, but are not consistent and may be an artefact of averaging low numbers of particles (Fig. S7).

Interestingly, while inspecting the nascent budding sites, we noticed an electron dense coat both on flat and negatively curved INM surfaces with cross-sectional dimensions similar to positively curved NEC (Fig. S8). This coat did not form a regular lattice, but rather an assembly of irregular rings approximately 25 nm in diameter (Fig. 4C, S8 b-e). These rings could be seen surrounding areas of regular lattice, suggesting that these structures are either a precursor or an intermediate form of NEC. Subvolume averaging of these rings did not result in a density map with a clear rotational symmetry, hinting at extensive heterogeneity and flexibility (Fig. S8g). Taken together, we hypothesise that the NEC transitions from a globally disordered layer, with some local sub-structure, to an ordered hexagonal array after reversible nucleocapsid binding. Once this transition takes place, envelopment proceeds regardless of the nature of the cargo.

### A fully assembled CVSC is not present in the nucleus or primary enveloped particles

As mentioned, it has been previously proposed that the CVSC, especially pUL25/17 (among others ^16, 48^), is responsible for mediating initial capsid-NEC interaction. Failing to determine any link between NEC symmetry and the capsid icosahedral orientation, we therefore decided to investigate whether there were any structures on the capsid that may indicate any difference in protein complement between perinuclear vesicles and nuclear or cytosolic capsids. We therefore obtained structures of the 12-fold vertices (pentons) to probe the molecular structure at this potential interaction site. We determined the molecular structure of PrV capsids contained within the nucleus, primary particles and the cytosol (Fig. 5, S9, S10). This allowed us to assign the composition of components of the capsid that were altered in their molecular arrangement during nuclear egress, as well as attribute any extra densities previously unidentified to potentially being significant to its mechanism.

**Figure 5.**
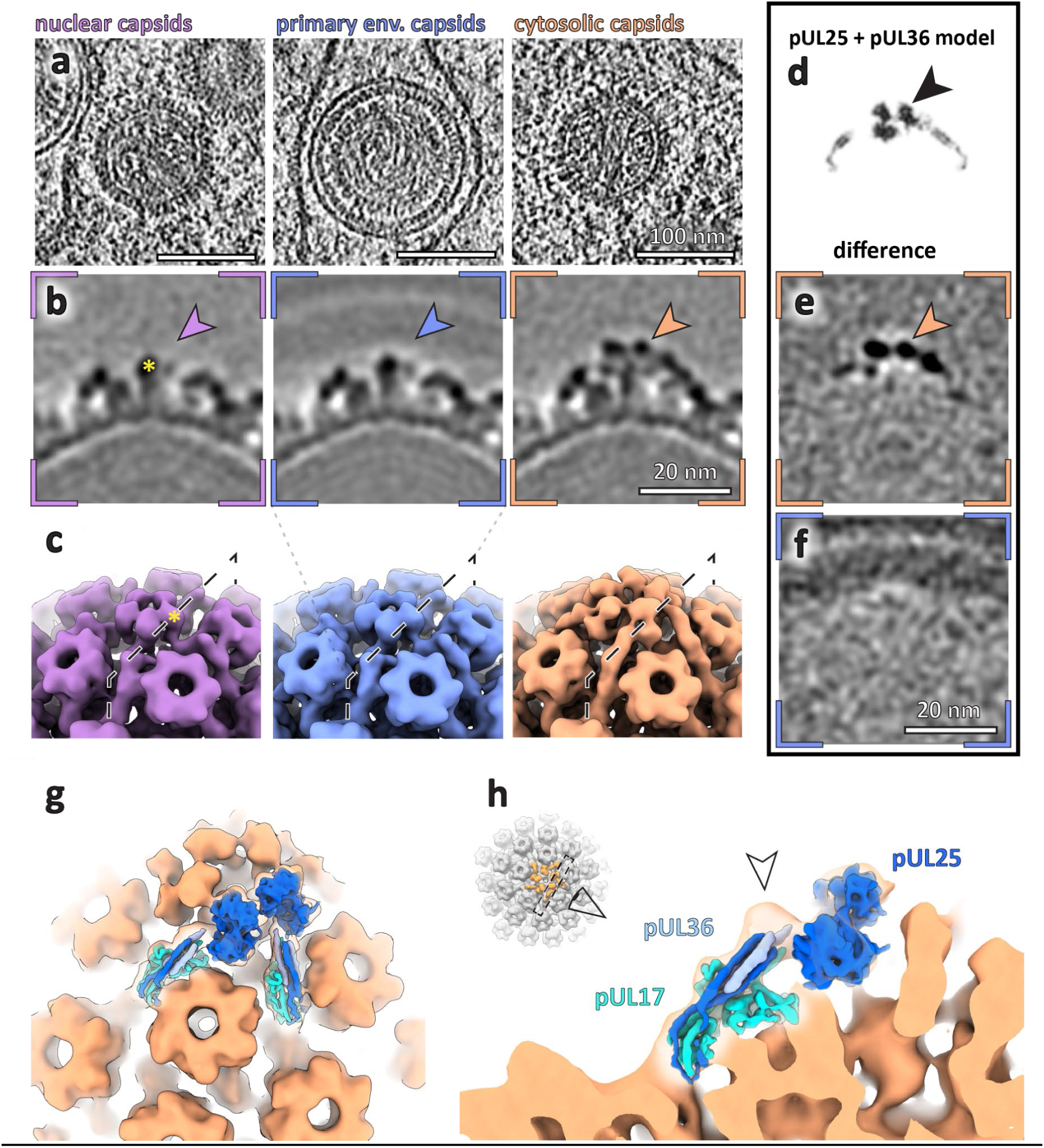
Subvolume averaging of PrV capsids in different subcellular locations reveals the compartment-dependent assembly of penton vertex components. a. Raw tomographic slices of capsids within specified locations within the cell. **b** Slices through the respective average volumes. Arrowheads indicate the position of a density present only in the cytosolic capsids. **c** Surface representation of volumes in b. **d** Slice through a simulated density map of two capsid vertex specific components (CVSC) pUL25 and pUL36 (PDB 2FSU, 7FJ1). **e** Normalised difference map between nuclear and cytosolic capsids. **f** Normalised difference map between nuclear and primary enveloped capsids. **g, h** Simulated density of CVSC components (PDB 2FSU, 7FJ1) fitted into cytosolic capsid map (orange). **h** Arrowhead indicates a density unaccounted for by existing atomic models (but present at low contour levels in e.g. EMD6387 and MD31611) .

The nucleocapsids contained within, and averaged from, the nucleus showed EM densities for pUL17, which has been previously attributed to the base of the CVSC found within capsids in fully-formed virions and was also suggested to be present in different numbers dependent on capsid type (A-, B- and C-type capsids)^49^; averaging the different types of capsids from within the nucleus did not exhibit major differences in structure at the resolutions we were able to achieve (S9, S10).

There was no significant difference between nuclear capsids and those determined within perinuclear vesicles at the resolution we achieved (24 Å, Fig. 5f). As expected, and consistent with our analysis of budding events, the NEC layer was smeared, with no obvious densities connecting the NEC layer and pUL17.

In contrast to nuclear and perinuclear capsids, A-, B-, and C- capsids in cytosol had a clear density for the full CVSC, previously shown to be contained in fully formed HSV1 and PrV capsids^18, 50^. The combined average of cytosolic capsids determined to ∼20 Å showed a good fit of both pUL17 and pUL25 (Fig. 5). The dimensions of reconstruction were consistent with the presence of the N-terminal portion of pUL36 at this resolution previously seen in single particle structures of virions ^49^. Difference mapping between the nuclear and primary particle capsid structures showed that in the regions of the reconstruction consistent with the 6-helix bundle (pUL17, pUL25 and pUL36 helices) and pUL25 globular domains to be the major structural difference between the capsids from the different subcellular regions. This implies that the CVSC (in its full form) is only formed once capsids have escaped from the nucleus and that there is no link between the NEC and the CVSC. The majority of cytosolic capsids used to determine the sub-volume average were in close proximity to the ONM, suggesting that they had escaped from the primary particles only recently. This implies that there is a component of the CVSC that is only recruited in cytosol.

pUS3 has been shown to be present on PrV capsids^51^, however, it is unclear that the deletion of pUS3 impacts upon the ability of pUL25 to associate with capsids in situ. Therefore, we attempted to determine whether the levels of recruitment of pUL25 in the nucleus is affected by deletion of pUS3. First, we verified that pUL25 is present at the same levels in WT PrV and the ΔUS3 mutant using fluorescence microscopy (Fig. S11). These results demonstrate that the recruitment of pUL25 is unaffected in our ΔUS3 system. Interestingly, fluorescent foci in the nucleus and cytoplasm had comparable intensity (both in WT and ΔUS3) and previous studies have also shown that pUL25 levels are unaffected by US3 deletion in vitro^52^. This might indicate that pUL25 is bound but requires a cytosolic component (perhaps pUL36) to become ordered. We also verified the CVSC assembly patterns using FIB/SEM and cryoET on WT PrV infected cells. Due to a very efficient nuclear egress, we observed no perinuclear vesicles, but the patterns of nuclear and cytosolic CVSC recruitment were identical to the ΔUS3 mutant. Similarly, procapsid assembly occurred in clusters or kindergartens despite the late stage of infection^53^ (Fig S12). Therefore, using PrV-ΔUS3 system reliably represents the processes that take place during WT PrV infection and CVSC assembles once capsids enter cytosol and it is not required for egress.

### Spheres, cylinders and planes

As mentioned previously, we used subvolume averaging in order to determine an electron microscopy density map of the NEC coat. The aim was to analyse the molecular architecture of the NEC and to further probe whether any inter-molecular interactions with capsids could be elucidated, or whether we could determine the molecular basis for curvature in these vesicles. We averaged all NEC-containing vesicles irrespective of size or shape. The final average reconstructed from ∼135,000 asymmetric units reached ∼ 14 Å resolution and shows a good match to previous data reconstructed from empty pUL31/34-GFP vesicles^10^ (Fig. 6, S13). The central hexameric ring was clearly separated into 6 sections, which we attributed to individual pUL31/34 dimers.

**Figure 6.**
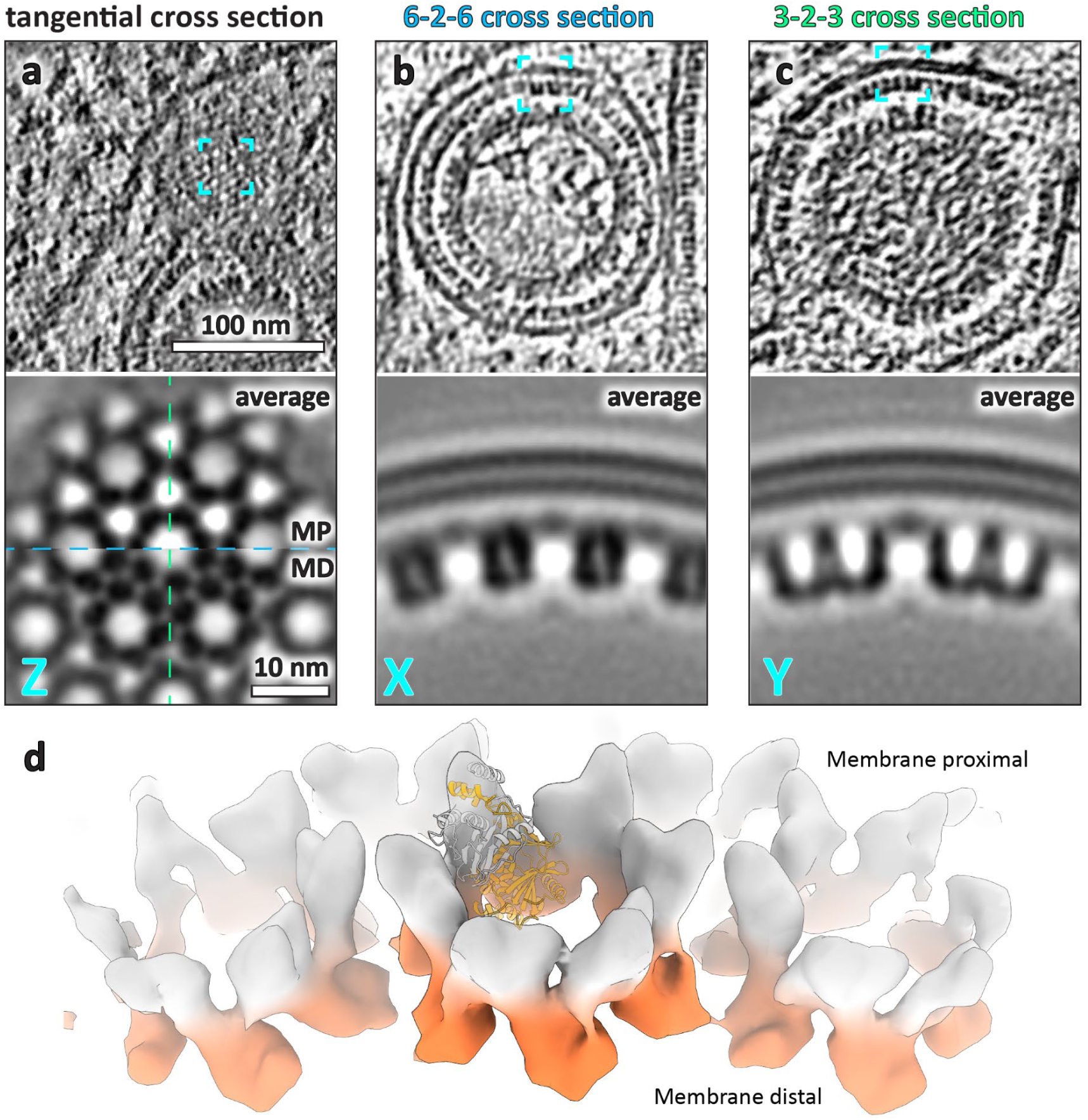
Subvolume averaging of PrV nuclear egress complex. **a** A tangential section through the NEC nuclear egress complex (NEC) within primary enveloped particles shown from the raw tomogram (top). Bottom row shows the density of the NEC subvolume average structure with the top half slicing through the membrane proximal layer and the bottom half through membrane distal layer. These represent the UL34 and UL31 parts of each heterodimer, respectively. **b** 6-2-6 or X direction and **c** 3-2-3 or Y direction of the NEC. **d** Isosurface representation of the average volume with the membrane proximal and distal layers highlighted in grey and orange, respectively.

In addition to the roughly spherical vesicles, we determined the subvolume averaging density map of the NEC in perinuclear tubes. The lattice on the inside of tubes was well ordered relative to perinuclear vesicles (Fig. 7). Small deformations did not correlate with a loss of long distance helical order, however, there were clear breaks in the lattice around bends (Fig. 7A). Notably, the inner diameter of observed tubes was between 60 and 72 nm, nearly half of spherical perinuclear vesicles and even empty vesicles reported previously^10^ (Fig. 7d, S2). The tubular average resolution was ∼21 Å despite being averaged from only ∼4,000 particles (c.f. for spherical average ∼135,000), suggesting that the tubular conformation may be more ordered/amenable to averaging. Notably, we were able to determine the structures of individual tubes with three different diameters and therefore three different helical parameters (Fig. 7e). This may indicate that there is no strong preference for the direction of curvature in the NEC lattice.

**Figure 7.**
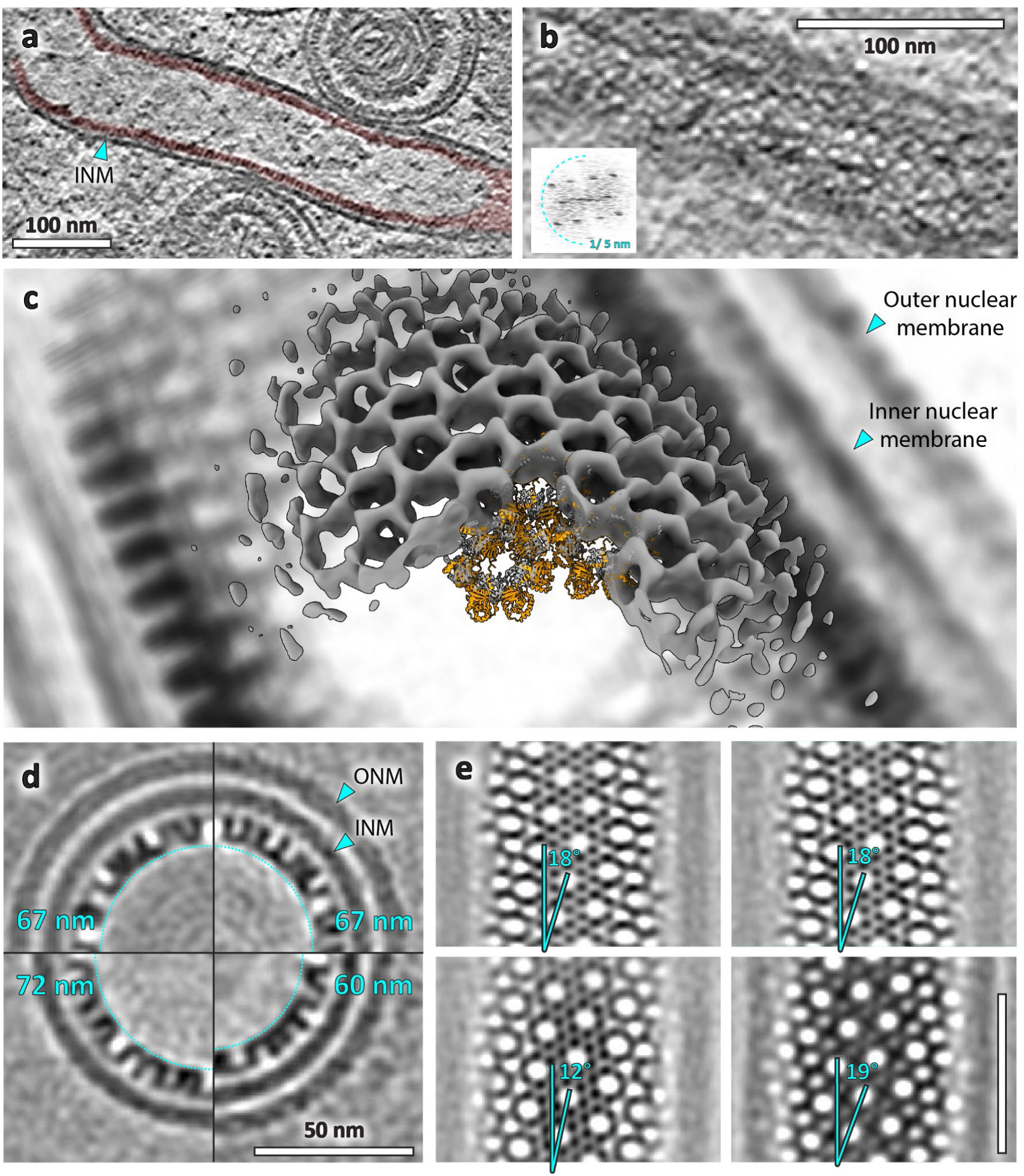
PrV nuclear egress complex forms tubes with helical symmetry. a. Slice through a tomogram showing a tubular NEC form (red) in type-1 NR. **b** tangential slice through the same tube showing a regular lattice arrangement. Inset shows the power spectrum of the projection through the lattice. **c** Isosurface representation of an average volume generated from 4 NEC tubes (overlaid with a density representation of a single tube). **d** Shown are sections through the average volumes of four tubes, their diameters are indicated in cyan. Observed NEC tubes have substantially smaller luminal cavities compared to canonical NEC vesicles. Notably, the two separate tubes with matching diameters and helical parameters were located within the same nucleoplasmic reticulum and may have originated from the same assembly. **e** Tubular NEC forms tubes with different helical parameters, indicating a flexibility in the direction of largest curvature. Highlighted is the angle of the 6-2-6 axis to the helical symmetry axis. Scale bar indicates 50 nm.

There was a third type of lattice in our data; a relatively flat double-layered sheet bringing together two surfaces of the INM. We postulate that this is the result of a head-head interaction between two NEC layers. Interestingly, a putative double-layer NEC observed previously in vitro consisted of individual rings reminiscent of the (single layer) rings we observed on negatively curved surfaces^40^ (Fig. S8). Considering that in the NEC hexamer found in perinuclear vesicles the distal portion of UL31 has an electrostatic potential that would repel UL31 in adjacent heterodimers, this configuration must require a conformational change that allows homotypic interaction with complementary electrostatics. This may explain the difference in subvolume average structures between these NEC formations, such as the hexamer-hexamer distance being substantially larger in the double-layer NEC compared to the spherical and helical lattices. Whether these non-canonical structures fulfil a specific function or are an aberrant state is not clear, but they might be useful for interrogating the flexibility of pUL31/34 heterodimers and of the NEC lattice.

### Fitting of the PrV pUL31/34 crystal structure into the NEC map suggests region of interest for curvature induction mechanisms

At the resolution of our subvolume average of the NEC lattice, a detailed all-atom analysis of conformational changes or interactions between and within individual heterodimers is not possible. However, attempting to fit the crystal structure of PrV pUL31/34 (PDB ID: 4Z3U) into the map does lead to some broad insights into the native, curved NEC structure. A simple rigid fit of multiple heterodimers shows that the lattice suggested by our map is consistent with the hexameric crystal lattice of the HSV1 NEC^22^ (Fig. 8A). Despite the obvious differences in long range curvature, the two lattices bear strong resemblances to one another, suggesting that the PrV NEC lattice structure from a previous study, modelled using a much lower resolution map and supported by analysis of point mutant functionality, is inaccurate^24^. The origins of the minor differences in the radius of the PrV hexamer (Fig. 8d, left and middle column) and the orientation and positioning of its heterodimers (Fig. 8b, c, left and middle column) as compared to HSV1 are difficult to ascertain, especially given the inherent flexibility in NEC conformations and configurations we have determined in this study. The relatively low resolution of the map could ascribe slightly erroneous positions for the fitting, or the differences might be inherent to the differences between HSV1 and PrV NEC lattices. The individual monomers in the hexameric layer may be able to adjust slightly to internal thermodynamic and structural changes imparted by the immediate surroundings of the NEC vesicle. It may also be created by mechanistic structural changes caused by change in curvature of the lattice, although the intra-hexamer changes seen do not seem capable of inducing curvature by themselves.

**Figure 8.**
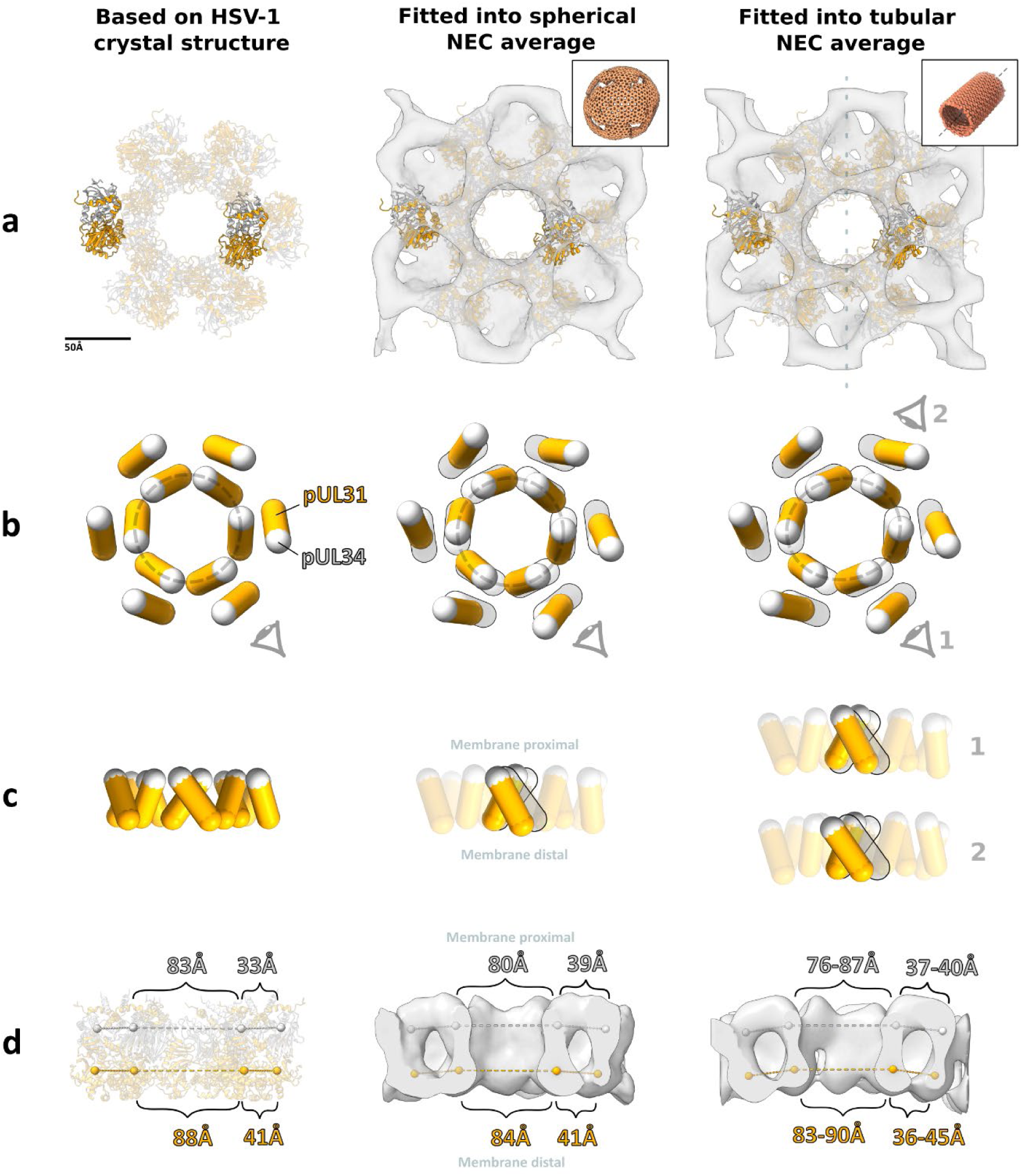
A comparison of different PrV NEC lattice structures, created by fitting PDB ID 4Z3U chains C/D into various structures. Left column, hexamer created by superposing PrV NEC (PDB 4Z3U) into heterodimer positions from crystal symmetry positions in HSV1 NEC (PDB 4ZXS), using ChimeraX *Matchmaker*. Middle column, heterodimer positions found by rigid body fitting of 4Z3U into subvolume average of the NEC from spherical/non-tubular vesicles. Right column, heterodimer positions found by rigid body fitting into subvolume average of NEC from tubular regions. **a** View of the three lattices with membrane proximal regions facing the viewer. Subvolume averages are shown as grey isosurface, pUL31 as an orange ribbon and pUL34 as a light grey ribbon. Dotted line in the tubular average (right) indicates the orientation of the helical symmetry axis. Insets depict overall lattice architecture from which the averages have been made. **b** Lozenge representation of heterodimers as seen in row a. Light grey markers represent centre of mass of pUL34 and arm of pUL31 (residues 18-55), orange markers represent centre of mass of the remainder of pUL31. Shadow lozenges in the middle and right columns represent the crystal structure-based model. Dotted grey ellipses show the shape of the central hexamer, as described by the centres of mass of pUL34/arm of pUL31. **c** Side view of lozenges as represented in b. View direction indicated by eye symbols in row b. **d** Distances between centre of mass markers on opposing sides of the NEC hexamers. Light grey and orange markers are as described in b. The tubular hexamer in the right column was elliptical, and hence the minimum and maximum distances are displayed.

A more significant change was found in the inter-hexamer distances (Fig. 8d, left and middle column). The membrane distal inter-hexamer distances of the curved PrV lattice and the flat NEC lattice are roughly the same, but the membrane proximal distances are larger in the PrV lattice by ∼6 Å. This increase is larger and more prominent than the intra-hexamer distance changes, and is in agreement with the type of movements expected to induce curvature. This suggests that interactions between hexamers at the membrane proximal region (most likely in pUL34) are responsible for curvature induction from the hypothetical flat lattice.

Notions that the PrV NEC hexamer remains a relatively inert unit, however, are broken by similar analysis of the heterodimers in the helical NEC average (Fig. 8, right column). Here, the intra-hexamer distances vary substantially, both from the previously analysed structures, and from themselves. The hexamer is stretched to form an ellipse, with the major axes 11 Å larger than the minor axes at the membrane proximal region, and 7 Å larger at the membrane distal region. Surprisingly, the major axes are not parallel with the direction of greatest curvature (i.e. perpendicular to the symmetry axis of the tube), which again suggests that intra-hexamer interactions are not a driving force for the induction of curvature.

### HSV1 nuclear egress follows the same patterns seen in PrV

Seeking to establish whether the patterns of nuclear egress we established are unique for PrV or are more broadly applicable to Alphaherpesvirinae, we performed FIB/SEM and cryoET on HSV1 infected cells (Fig. 9). Similarly to WT PrV in the natural host cells, we did not find any nuclear egress events, though we were able to identify ring-like NEC structures on the INM resembling those found in the ΔUS3 PrV system (c.f. Fig 9, S3). In Vero (green African monkey) cells, however, we captured 4 egressing capsids and one event of NEC budding (Fig. 9a) and could therefore confirm the patterns of CVSC assembly observed in PrV: pUL17 is present in nuclear and perinuclear capsids, but pUL25 is only assembled (or becomes ordered) in the cytoplasm (Fig 9c). We also noticed examples of non-canonical events, such as a primary enveloped B-capsid and double-layer NEC. Subvolume averaging of ∼9,000 particles of the HSV1 NEC from perinuclear vesicles resulted in a map with ∼21 Å resolution (Fig. 9b, S14). The map is similar to that of the PrV NEC with subtle differences. The most notable difference is the orientation of the dimer subunits, visualised by comparison of rigid-body fitted atomic models. While in PrV the pUL31/34 dimer is tilted by ∼35° to the six-fold axis, the HSV dimer is tilted by ∼25° (Fig. S14c). As a result, the curved HSV NEC hexamer is very similar to the in vitro crystal lattice of soluble pUL31/34. Assuming that the crystal lattice is a reliable approximation of a flat lattice, this could further support the notion that it is the inter-hexamer interfaces that are the direct cause of curvature formation. At first glance, it is difficult to reconcile the large difference between dimer orientations in HSV and PrV with a (presumably) common curvature inducing mechanism. Nevertheless, the high degree of flexibility within the lattice observed in PrV could also allow for a gradual accumulation of mutations and subsequently divergence within the hexamer.

**Figure 9.**
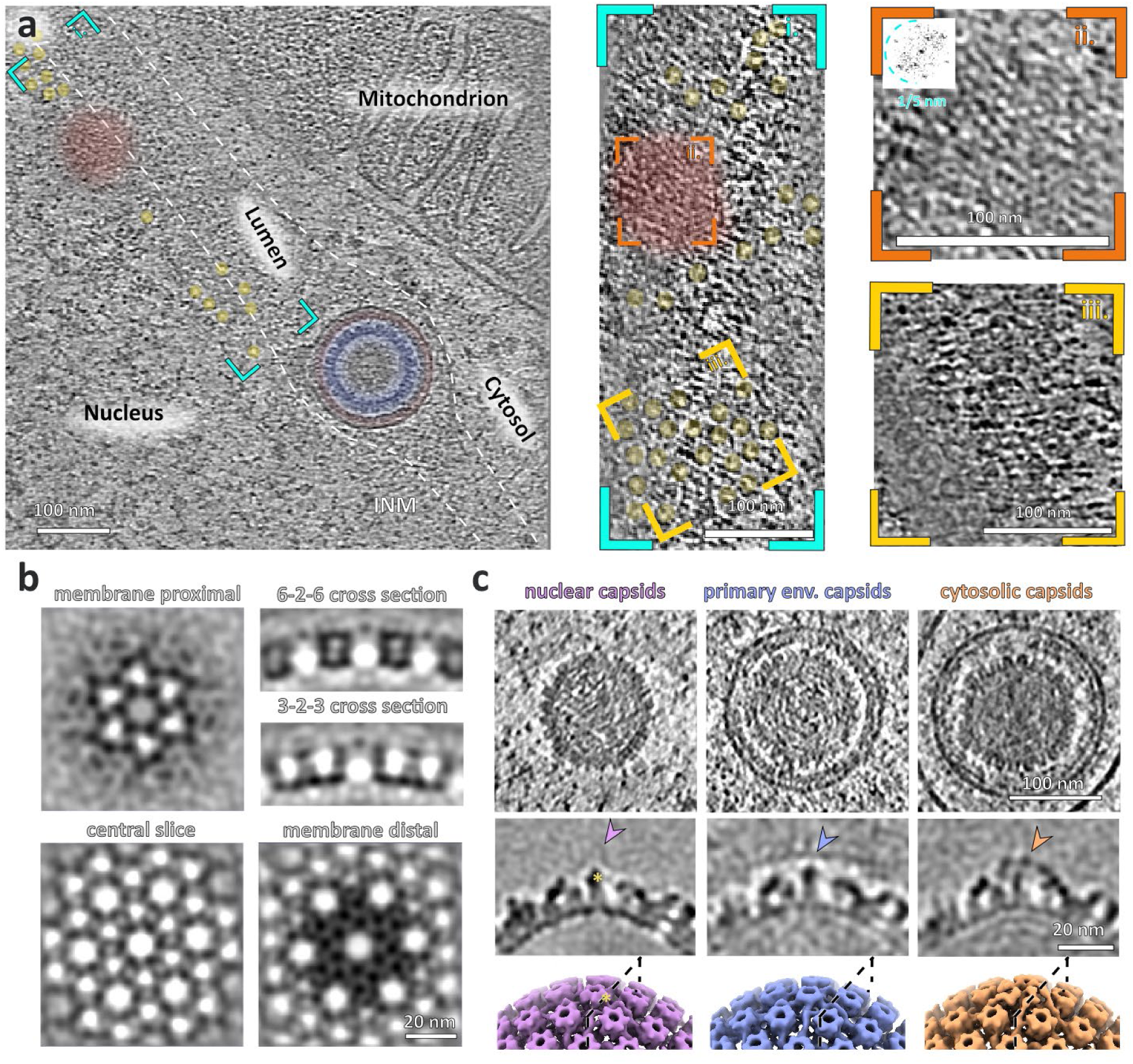
NEC in HSV1. **a** Slice through a tomogram depicting primary enveloped nucleocapsid egressing to the cytosol. The NEC coat of WT HSV1 (red) follows the same pattern as that of ΔUS3 PrV and could be identified on the INM and in the perinuclear vesicles. Likewise ring-like structures similar to those in PrV (yellow in i. and iii., Fig 4c and S8) was identified near the hexagonal NEC lattice (red in i. and iii.). Red area in the insert ii. shows budding NEC. **b** HSV1 NEC crystal structure (PDB ID 4ZXS) fitted into the subvolume average of HSV1 NEC from perinuclear vesicles. **c** Raw tomographic slices through C-capsids of WT HSV1 in indicated subcellular locations. **d** Slices through the average volumes of C-capsids (middle) and their surface representations (bottom). Arrowheads indicate the position of an additional density present in the cytosolic capsids.

In summary we show that in situ, the NEC modulates the INM and ONM in many configurations both in the presence and absence of nucleocapsids. This inherent flexibility is explained by the lack of pentamer (five-fold) organisation, where to produce perinuclear vesicles the NEC hexamer must be deformable. Our subvolume averaged lattices of NEC perinuclear vesicles and tubules demonstrate that there is substantial freedom of movement between heterodimers within and between hexamers. In other words, the NEC lattice is remarkably plastic, which is likely required to compensate for the deformations caused by spherical curvature applied to a hexameric lattice. Our observations lead to the conclusion that NEC-capsid recruitment is not a rigid process, with capsid proximity not necessarily being a driver of NEC formation and elongation. The proximity of a capsid regulates the size of the forming vesicles, however, this is not mediated specifically by proteins at the five-fold vertex of the capsid but rather by the electrostatic interactions between the distal region of pUL34 and the VP5 capsid protein. These data are consistent with previous observations that herpesvirus egress is similar to RNP aggregate removal from the nucleus^5, 10^, insofar as pUL34/31 heterodimers can associate to form intraluminal arrangements diverse in both size and geometry. Further work will be needed to understand whether there is a role for these arrangements in the herpesvirus life cycle or if they are an evolutionary redundant pathway relating to molecular remodelling of nuclear membranes.

## Acknowledgements

We gratefully acknowledge funding by the Deutsche Forschungsgemeinschaft (DFG) under Germany’s Excellence Strategy EXC 2155— project no. 390874280 and by the Wellcome Trust through a Collaborative Award (209250/Z/17/Z) as well as Hamburg-X and the the Deutsche Forschungsgemeinschaft (DFG, German Research Foundation) - GRK2771 – project no. 453548970 (J.B., K.G.), Wellcome Trust grants 107806/Z/15/Z and 209250/Z/17/ Z (KG), BMBF grant 05K18BHA (KG), Wellcome Trust grants 099683/Z/12/Z and 225902/Z/22/Z (M.G.). This research was funded in part by DFG INST 152/ 772-1, 774-1, 775-1, 777-1 FUGG (CSSB cryoEM facility) and a Wellcome Trust core award to The Wellcome Centre for Human Genetics (090532/Z/09/Z). V.P. was supported by a Nuffield Dept of Medicine Prize Studentship.

## Materials and Methods

### Cell Culture

#### PrV-ΔUS3

A mutant PrV-ΔUS3 virus ^54^ containing a GFP-positive selection marker was used to infect immortalised pig-epithelial cells (EFN-R; CCLV-RIE 0089) grown on gold, holey-carbon coated electron microscopy grids and vitrified using a manual plunger with liquid ethane propane mixture (37%/63%).

#### WT PrV and HSV1

EFN-R cells (CCLV-RIE 86), Vero cells (ATCC CCL-81) and immortalised HFF cells (ATCC CRL-4001) were grown at 37°C with 5% CO_2_ in DMEM medium supplemented with 10% FBS and 1% Glutamax (Gibco). Gold grids with SiO_2_ film (Quantifoil, Au 200 mesh, R 1/4) were glow-discharged and coated with 20 µg/ml fibronectin solution for 30 min. Trypsinized cells were applied to grids and incubated for a minimum of 4h and up to 12h. WT PrV (Kaplan) was added to the cells at MOI=30 and incubated for 12h. Separately, WT HSV1 (strain 17) was added to the cells at MOI=30 and incubated for 12-18h. All samples were plunge-frozen either on Leica GP2 plunger or manual plunger in liquid ethane propane mixture (37%/63%).

### Lamellae Production Dual-Beam Scanning Electron FIB-SEM Cryo-Microscope

#### PrV-ΔUS3

Lamella were milled using a dual-beam focused ion beam/scanning electron “Scios” microscope (Field Emission Industries, Inc, now Thermo Fisher Scientific, Oregon, USA) equipped with a Quorum PP3010 cryo transfer system. Milling was performed as outlined^55^. Data were acquired from a total of 29 lamella.

To probe the NEC-capsid interactions during herpesvirus nuclear egress, we targeted regions on the lamellae that contained enveloped nucleocapsids located within the lumen between the INM and ONM. At intermediate magnification (5,600x, nominal) it was possible to identify individual primary vesicles and discern the characteristic electron-dense coat in between the nucleocapsid and membrane; these features were used to target tilt series collection at 61,000 x nominal magnification.

#### WT PrV and HSV

Vitrified HSV1 and WT PrV infected samples were clipped in autogrids modified for FIB-milling^55^ and transferred to Aquilos focused-ion beam scanning microscope (Thermo Fisher Scientific). Lamellae of approximate thickness between 100-250 nm were prepared using automated procedure with AutoTEM software (Thermo Fisher Scientific).

### Electron Cryo-Tomography Data Acquisition

#### PrV-ΔUS3

Electron cryo-tomography data were acquired on lamella using a Tecnai G2 Polara transmission electron microscope (FEI) equipped with a field emission gun operated at 300kV, a GIF 2002 post-column energy filter (Gatan, Pleasanton, CA), and a Gatan K2 Summit Direct Electron Detector (Gatan, Pleasanton, CA) operating in counted mode. Tomographic tilt-series acquisition was performed under low-dose condition with a total of 30 tilt series from 27 lamellae. The tilt range used was from -60 to +60, with cumulative doses varying between ∼ 80 – 200 e^-^/Å^2^. Data acquisition was performed using SerialEM^56^, with tilt-series acquired in a bi-directional manner using 3° tilt increments, a defocus range of 3-6 µm defocus and at a calibrated pixel size of 3.55Å/px.

#### WT PrV and HSV

Vitrified lamellae of WT HSV1 or WT PrV infected cells were imaged on a Thermo Fisher Scientific Titan Krios TEM operating at 300 kV, equipped with a field emission gun (XFEG) and a Gatan Bioquantum energy filter with a slit of 20 eV and a Gatan K3 electron detector. Tilt series were acquired using SerialEM software with cumulative dose of 100-120 e^-^/Å^2^, tilt range of -52 to +67 with the starting angle of 8° in dose symmetric manner^57^ and tilt increment of 3°. Defocus used ranged between 4-6 µm. Data was acquired at pixel sizes of either 2.15 Å/px for WT HSV1 or 3.336 Å/px for WT PrV.

### Sub-Volume Averaging

Tomograms were reconstructed using IMOD Etomo or Aretomo^58^. WT PrV tilt series were scaled to pixel size 3.55 Å/px, CTF corrected using IMOD ctfplotter and then reconstructed in Aretomo^58^.

#### Canonical NEC

Sub-volume averaging was performed largely similar to previously described for NEC vesicles in the absence of viral capsid ^10^. Briefly, an initial reference was extracted from the tomogram which was centred on a single hexon position within a perinuclear vesicle. This raw sub- volume was then used as the initial reference. 20 initial vesicles were modelled as a perfect sphere with a radius which was manually determined to be the rough diameter of a given vesicle in IMOD^59^. The model positions were sampled so that one point was roughly the space of an inter-heterodimer distance (10 nm). Subsequently, the model points were allowed to move in an iterative fashion using the sub-volume averaging software PEET^60^ with the sampling becoming progressively finer. Initial sub-volume averaging was performed on rescaled data until no improvement in resolution was observed, and then the determined results applied to the original sampling, with six-fold symmetry applied, until the structure converged. This average was then used as a reference for the alignment of a further 37 vesicles modelled similarly as spheres, and 68 irregularly shaped vesicles and membrane patches containing NEC, which were picked using IMOD segmentation tools from 16 tomograms. These particles were aligned in a similar fashion to the original 20 vesicles, starting with data binned four times and gradually binning less and removing badly positioned, overlapping and poor scoring particles across five PEET runs. Six-fold symmetry was applied by creating duplicate particles with orientations rotated around each particle’s symmetry axis. A total of 22,596 particles were symmetrised, so that 135,576 particles were used in the final average. The FSC for the NEC averages were calculated by splitting the final dataset in half, randomising the orientations of all particles and realigning separately. FSC plots were generated using Bsoft^61^. A soft-edge spherical mask (85 nm diameter) was applied to both half-maps. All sub-volume averaging was performed using the “gold-standard” method and the resolution was taken at the 0.143 criterion. For HSV-1 NEC vesicles the same procedure was followed as described above using a canonical PrV NEC map as an initial reference binned 4 times. 4 spherically shaped vesicles from 4 different tomograms were available for particle picking. A total of 1421 particles were used, resulting in 8526 particles in the final average after symmetrisation.

#### Nascent NEC

Model points were picked manually at three sites containing capsids in proximity to NEC patches, and their Y axes (normal to the membrane) were oriented to the nearest capsid centre. An initial reference was generated by averaging manually picked positions with random rotations around the Y axis. The average volume converged into a hexagonal lattice resembling the canonical NEC map. Subsequently, particles from 28 budding events generated using IMOD drawing tools were aligned to an average volume of canonical NEC from perinuclear vesicles with volumes binned 4 times. Aligned particles were first cleaned by cross-correlation coefficient, and then manually cleaned to remove those that were outside hexagonally arrayed regions or those that were clearly misaligned (e.g. centred on the membrane rather than the NEC layer). In order to determine the positions of particles missed by manual picking, the lattice formed by the particles was expanded to create new model points, by assuming C6 symmetry and an interhexamer distance of 110 nm, then aligned using a cylindrical mask with 22 nm diameter (roughly a single hexamer), and again cleaned manually and by cross-correlation coefficient. This process was repeated until the number of points remained roughly constant. The resolution of the final map was not determined.

#### Tubular NEC

The 4 tubular NEC structures were first averaged independently. Initial references were obtained by averaging a small number of particles aligned to the cylinder symmetry axis and random rotations. These independently converged to a recognisable NEC lattice structure. Particle coverage was expanded from this initial particle seed using manually determined helical symmetry (-18.5°, -17°, and -12.16° twist and 22.7 Å, 22.7Å, and 71 Å rise, respectively). The 4 independent particle sets were then combined and averaged with progressively less binning. C2 symmetry was applied by adding particles rotated by 180° perpendicular to the helical symmetry axis. For FSC determination, particles were split after alignment with 2 times binned volumes and their rotations randomised by up to 5° along a randomly selected axis.

The two halves were then aligned to the respective references obtained by particles with the randomised orientations.

#### Nucleocapsids

EMD-6907 was used as an initial reference with volumes binned 8 times (57 Å Nyquist), followed by alignment of particles extracted at penton positions. These particles were split into two halves. Symmetry was applied by adding particles rotated by multiples of 72°. Alignment and averaging was performed with volumes binned 4 and 2 times and finally with unbinned volumes (3.55 Å Nyquist). The two half datasets were then combined, aligned, and filtered to their respective resolution at 0.143 FSC cutoff using Bsoft and an arbitrarily chosen B factor.

### Data visualisation

Volumes were rendered using IMOD, Chimera, ChimeraX or Matplotlib^62–64^. Manual segmentation of membranes was done manually using IMOD tools. Backplotting, the placement of average volumes into the original tomogram coordinate system using positions and orientations determined by subvolume averaging was performed using Python scripts based on TEMPy ^65^. Segmentation was done manually using IMOD drawing tools.

### Rigid body fitting

In order to create pseudo-atomic models of the spherical and tubular NEC lattices, the crystal structure of PrV UL31/34 (PDB 4Z3U) was fitted as rigid bodies into the relevant maps. An initial hexamer, together with six interacting heterodimers from neighbouring hexamers, were built by superposing the PrV structure atop the positions found in the symmetry expanded HSV-1 UL31/34 crystal structure (PDB 4ZXS). The ChimeraX *Matchmaker* program was used for this superposition.

These twelve heterodimers were then fitted together as a single rigid body into the two NEC density maps, using ChimeraX *Fit in Map*. Each individual heterodimer was then fitted individually with the same program, using the previous fit as an initial position. Marker positions for the centre of mass of UL31 residues 56-271 and UL34 + UL31 residues 18-55 were calculated using the ChimeraX *measure* command, and the inter-marker distances using the distance dialogue box.

### Curvature analysis

Curvature analysis was performed by fitting a central surface normal and 2 principal curvatures to each central hexon and its neighbours within 426Å (60 voxels x 7.1Å voxel size) using scipy.optimize^66^. Random perturbation of initial estimates and leave-one-out cross-validation were used to avoid overfitting and issues with local optima. This process was run for all subvolume particles from canonical NEC vesicles, and from one representative tubular NEC lattice.

### Fluorescence microscopy

#### Cell culture

PK15 cells (CCL-33) were cultivated in Dulbecco’s modified Eagle’s medium (Gibco) containing 10 % (v/v) foetal calf serum (Gibco) and kept at standard cell culture conditions (37° C, 95 % relative humidity, and 5 % CO2).

#### Viruses

PrV-mScarlet-UL25 was made by homologous recombination by transfecting PK15 cells with phenol-chloroform extracted WT PrV DNA (strain Kaplan^67^) and an 893 bp PCR-fragment coding for an insertion of mScarlet-I ^68^ 5’ between amino acids 42 and 43 of UL25 as previously reported^69^. PrV-mScarlet-UL25-ΔUS3 was generated similarly by homologous recombination through transfecting PK15 cells with phenol-chloroform extracted PrV DNA strain Kaplan ΔUS3^54^ but using a synthetic DNA construct (Biomatik, Canada) employing longer homologous sequences of 399 bp upstream and 573 bp downstream. After recombination, fluorescent viral clones were selected and underwent three rounds of plaque purification on PK15 cells.

#### Viral infections and sample preparation for microscopy

PK15 cells were seeded on 35 mm dishes (ibidi) and infected at a multiplicity of infection (MOI) of approximately 50, resulting in a mostly synchronised infection of all cells. After 1 h, the virus-containing media was removed, cells were washed with 1 mL of PBS, and new media was added. At indicated time points, cells were fixed for 15 min at room temperature with paraformaldehyde (4%) in PBS and washed three times with 1 mL of PBS. To counterstain nuclei, cells were incubated with Hoechst 33342 (Thermofisher) diluted to 16 µM in PBS and washed with PBS after 15 min.

#### Spinning disc fluorescence microscopy

Volumes of infected cells were acquired with a Nikon Eclipse Ti2 equipped with a Yokogawa CSU-W1, an Andor iXon Ultra DU-888U3 EMCCD camera, and an SR Apo TIRF AC 100xH objective (NA = 1.49) using 405 nm and 561 nm lasers with a quad filter (405/488/568/647) and 447/60 as well as 600/25 emission filters at a step size of 200 nm.

#### Single particle detection

Viral particle intensities were measured using Trackmate v6^70^ in FIJI^71^. Nuclear and cytoplasmic areas were marked as regions of interest (ROI), the expected blob diameter set to 0.4 µM, and the quality threshold set to 10.000. Total intensities were extracted from the resulting XML files and quantified with Origin.

## Notes

### Competing Interest Statement

The authors have declared no competing interest.

