## Supplemental information for "NECing goes: flexibility of the herpesvirus nuclear egress complex"

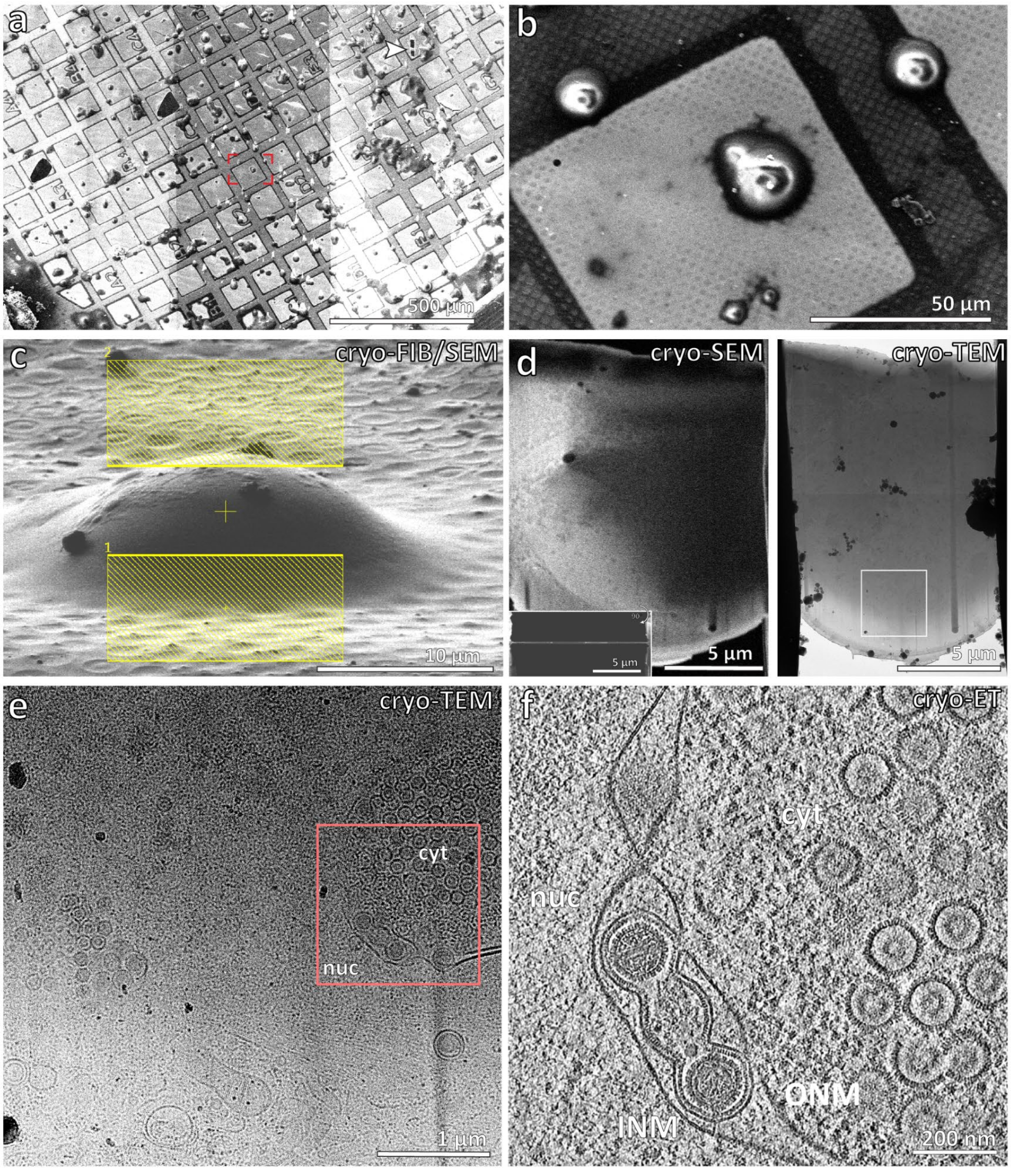


**Figure S1. Workflow for CryoFIB/SEM and CryoET.** **a** A low magnification SEM image of PrV-ΔUS3-infected porcine epithelial cells. **b** a Higher magnification image of the box shown in a. **c** An oblique SEM view illuminated by the FIB with yellow boxes indicating the area above and below the cell for targeting with the focussed ion beam. **d** SEM image of thinned cellular section (lamella) from the top and side (inset). Transmission electron microscope (TEM) image is also shown (right), with target area shown in e highlighted by a white box. **e** A TEM image shown at 9500x nominal magnification. Details of the cell are visible at this magnification, allowing targeting of regions of interest for higher magnification tomographic data collection (red box). cyt = cytosol, nuc = nucleus. **f** A tomographic slice of the region indicated in e.) at 35000x magnification, nominal.

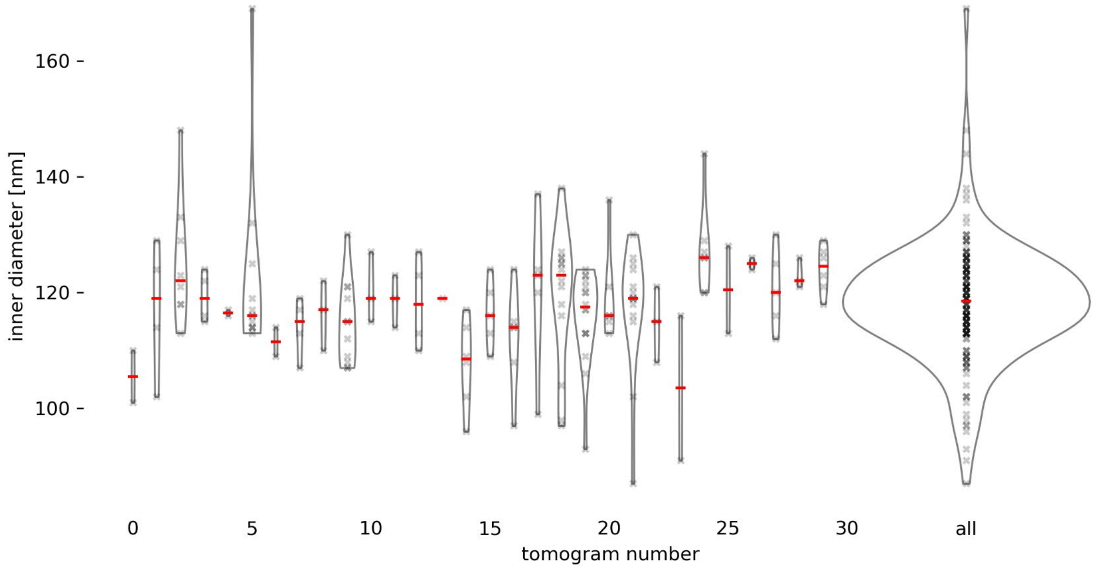


**Figure S2. Sizes of observed near-spherical perinuclear particles.** Shown are inner luminal dimensions of vesicles in each tomogram, the width of violin plots is proportional to the number of measurements. Red lines indicate medians. Vesicles were measured manually by centering model points on membrane density on opposite sides of periplasmic vesicles. To get the inner diameter, 34 nm (the approximate thickness of two membranes and NEC layers) was subtracted. Diameters smaller than 120 correspond to empty vesicles.

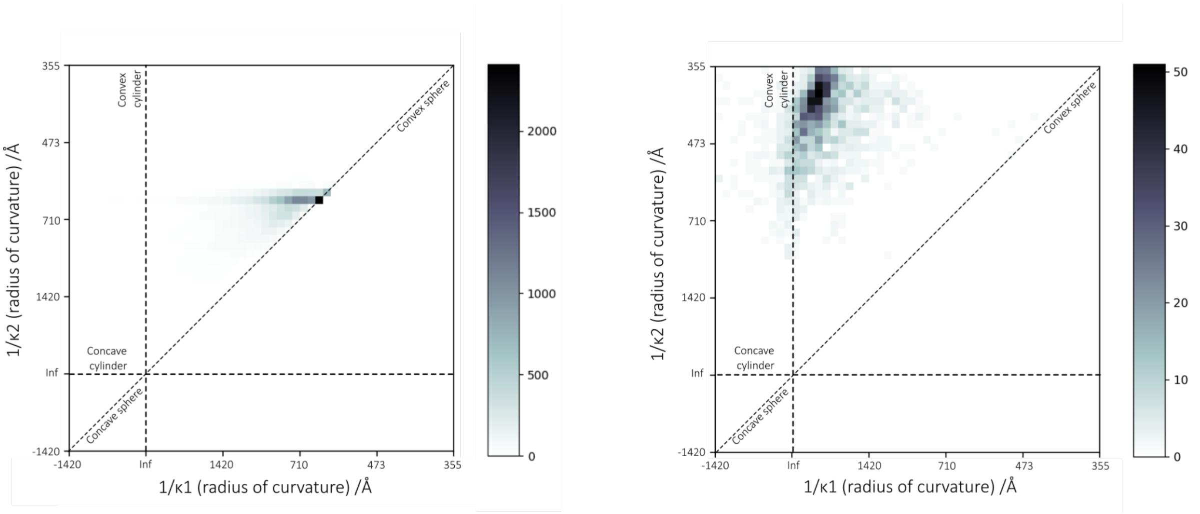


**Figure S4. Curvature analysis of PrV NEC.** 2D histograms of radii of curvature of NEC particles from canonical vesicles (left) and tubular vesicles (right). The diagonal dotted line in each represents the case where both perpendicular radii of curvature are identical, thus describing a sphere. The Inf/Inf bin of each histogram describes infinite radii of curvature in both directions, thus describing a flat plane. The vertical and horizontal dotted lines represent an infinite radius of curvature in one direction, thus describing a cylindrical surface. In this case, a negative radius of curvature represents a concave NEC surface (ie, with the NEC on the outside of a spherical vesicle). The histogram peak for the tubular NEC is ~39nm, and for the canonical NEC it is ~62nm. Remaining canonical particles tend toward ellipsoidal curvature rather than larger spherical topology.

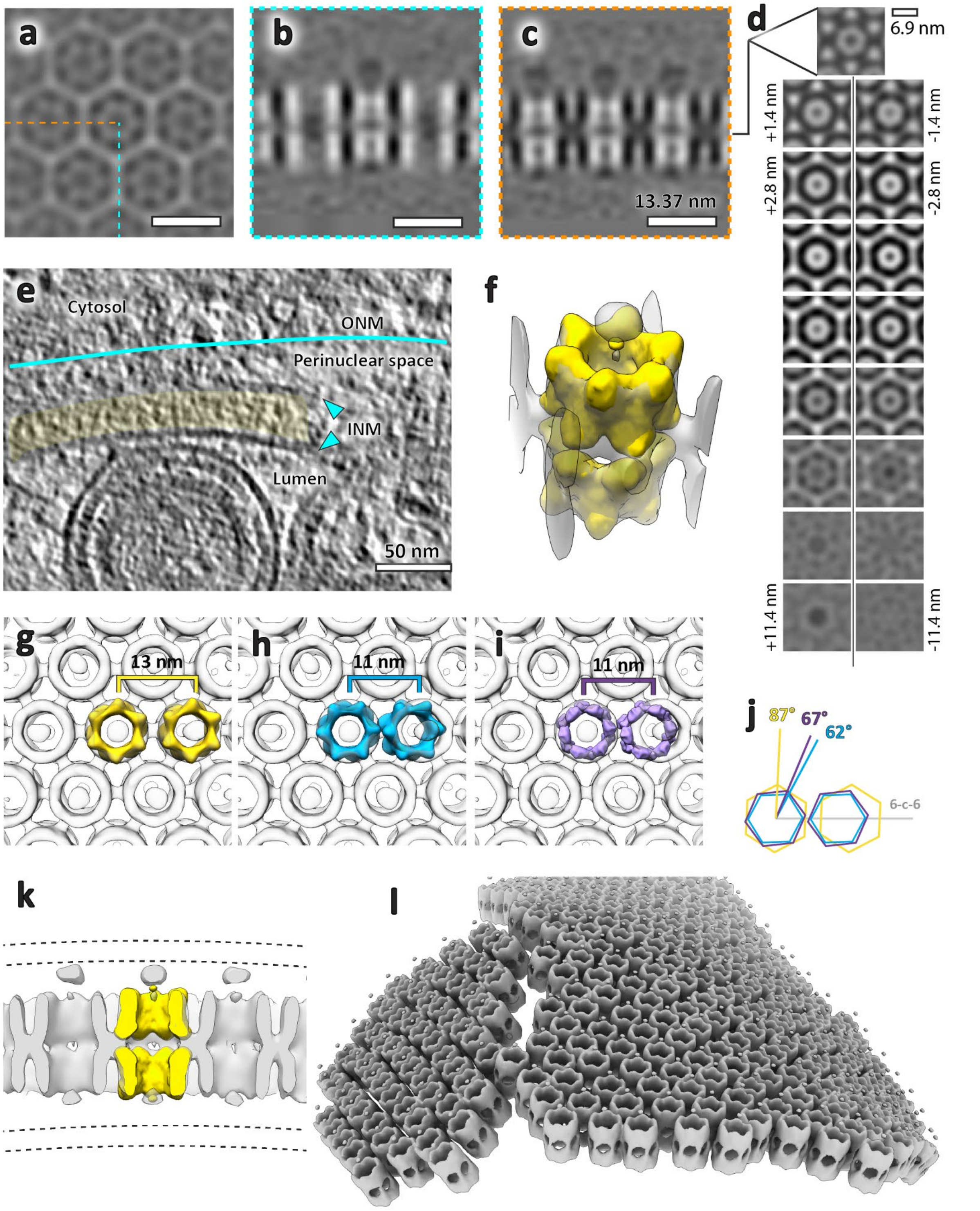


**Figure S3. Putative flat NEC double lattice. a, b, c** Orthogonal section through average volume of lattice shown in Fig. 2f. There is no density for the membrane bilayers due to the majority of particles having their six-fold symmetry aligned with the tomogram Z axis. A total of 1944 C6 symmetrised particles were included in the average. **d** Tangential sections through the volume at 1.4 nm intervals. **e** A slice through the raw tomogram indicating the position of the inner nuclear membranes relative to the lattice. The membrane is not visible around the majority of the lattice layer due to its orientation to the tomogram missing wedge. The approximate position of the outer nuclear membrane was inferred from the exclusion zone of cytosolic components (ribosomes, intermediate filaments, microtubules). **f** Two NEC hexamers segmented from the canonical lattice (Fig. 7, yellow) were fitted into one repeating unit of the lattice. In this orientation, pUL31 would form the interface between the two lattice layers. **g, h, i, j** The hexamer centres in the double lattice (yellow) are spaced

2 nm further apart and are rotated by approximately 20° to the 6-2-6 axis compared to the canonical NEC (blue) and flat lattice derived from the HSV1 crystal structure (purple). **k** The position of membrane bilayers was estimated from the raw tomogram and comparison to canonical NEC. **l** Plotback of individual symmetry units (dodecamers). There is a slight curvature to the lattice with a break in the middle, presumably to accommodate the tighter curvature of the underlying nucleoplasmic reticulum membrane.

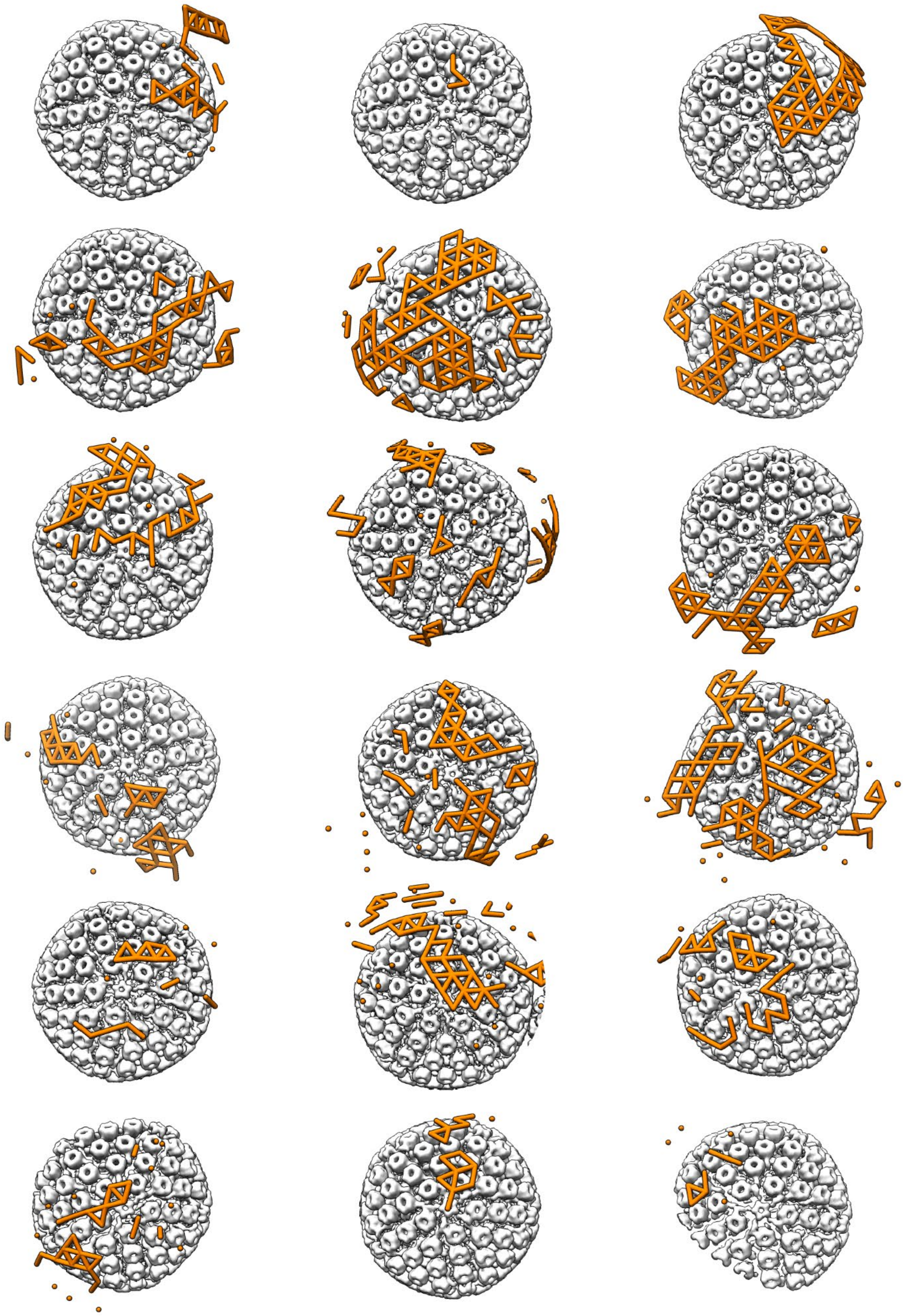

**Figure S5. 3D overview of putative budding events.** Capsid volumes are aligned with one penton vertex facing the viewer. Orange bars connect NEC hexamer centres separated by less than 12 nm.

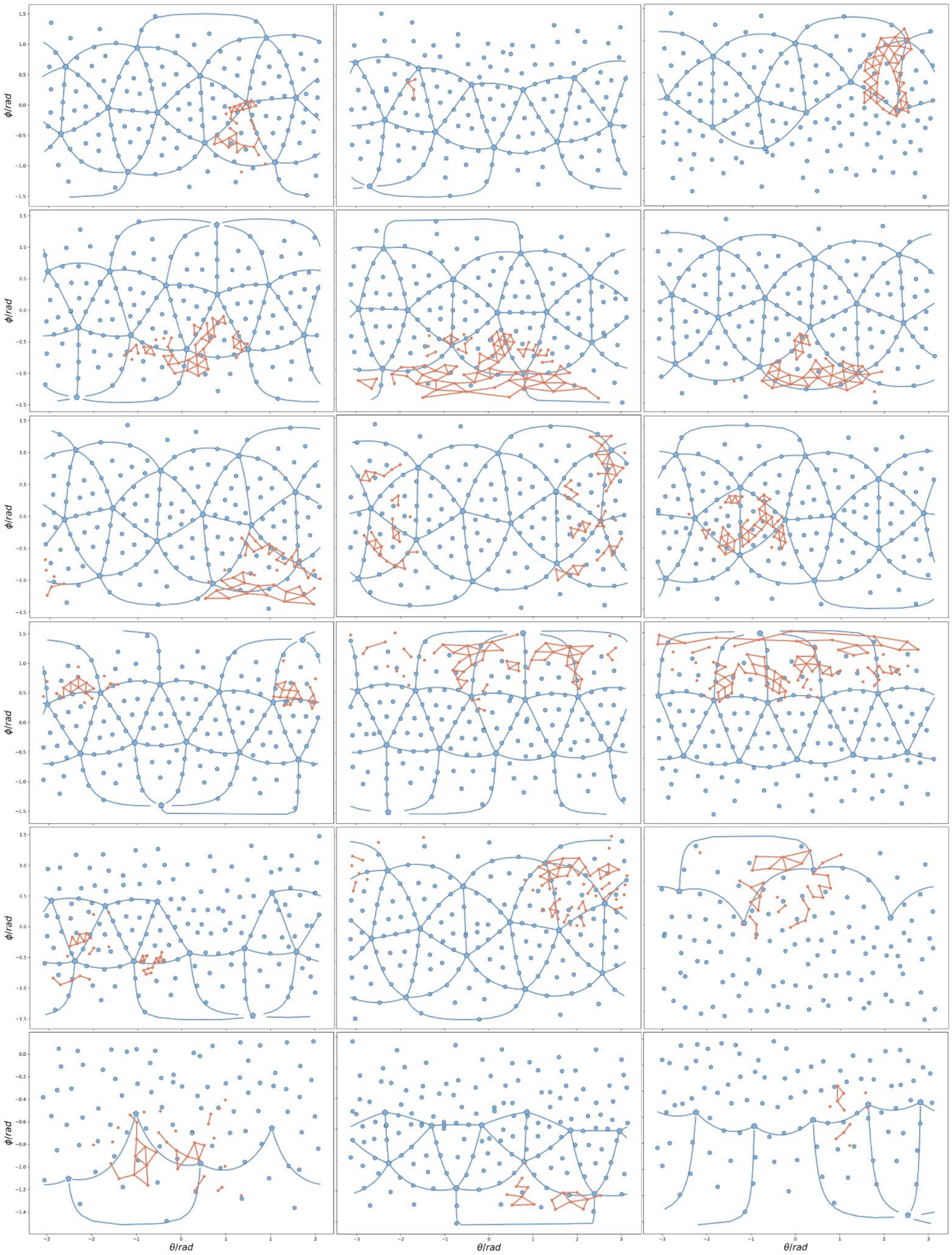


**Figure S6. Radial coordinate overview of putative budding events**. These events are the same as those shown in Fig. S5, but represented in radial coordinate space with the capsid centre as the origin. Blue lines represent icosahedral edges, pentagons represent penton vertices, and circles represent hexons. Some hexon and penton positions are obviously misaligned, this is due to some capsids being only partially contained within the lamella.

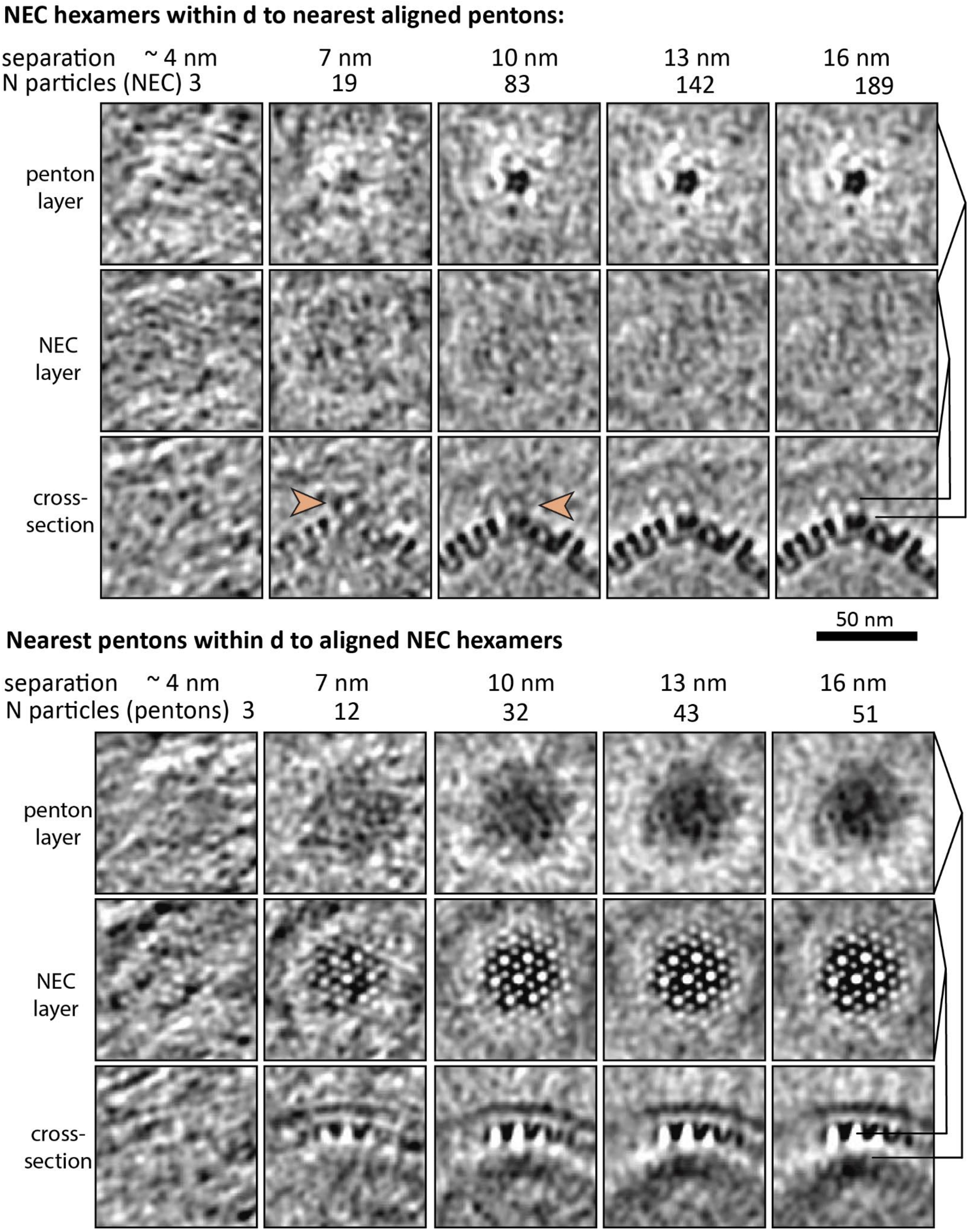


**Figure S7. Questionable connecting densities between nuclear capsids and budding NEC.** NEC were classified by the distance d, to the nearest penton vertex (top panel) and vice versa (bottom panel). Each column shows sections through the resulting class average volume, with the maximum separation distance and the number of particles included in the average indicated above. All volumes were filtered using bfilter bandpass filter. The NEC layer is smeared in the penton-aligned averages and accordingly the capsids are smeared in the NEC-aligned averages, indicating that the two lattices are not aligned. However, there is a hint of a density originating from pentons closer than ~7 nm from the nearest NEC surface (orange arrows).

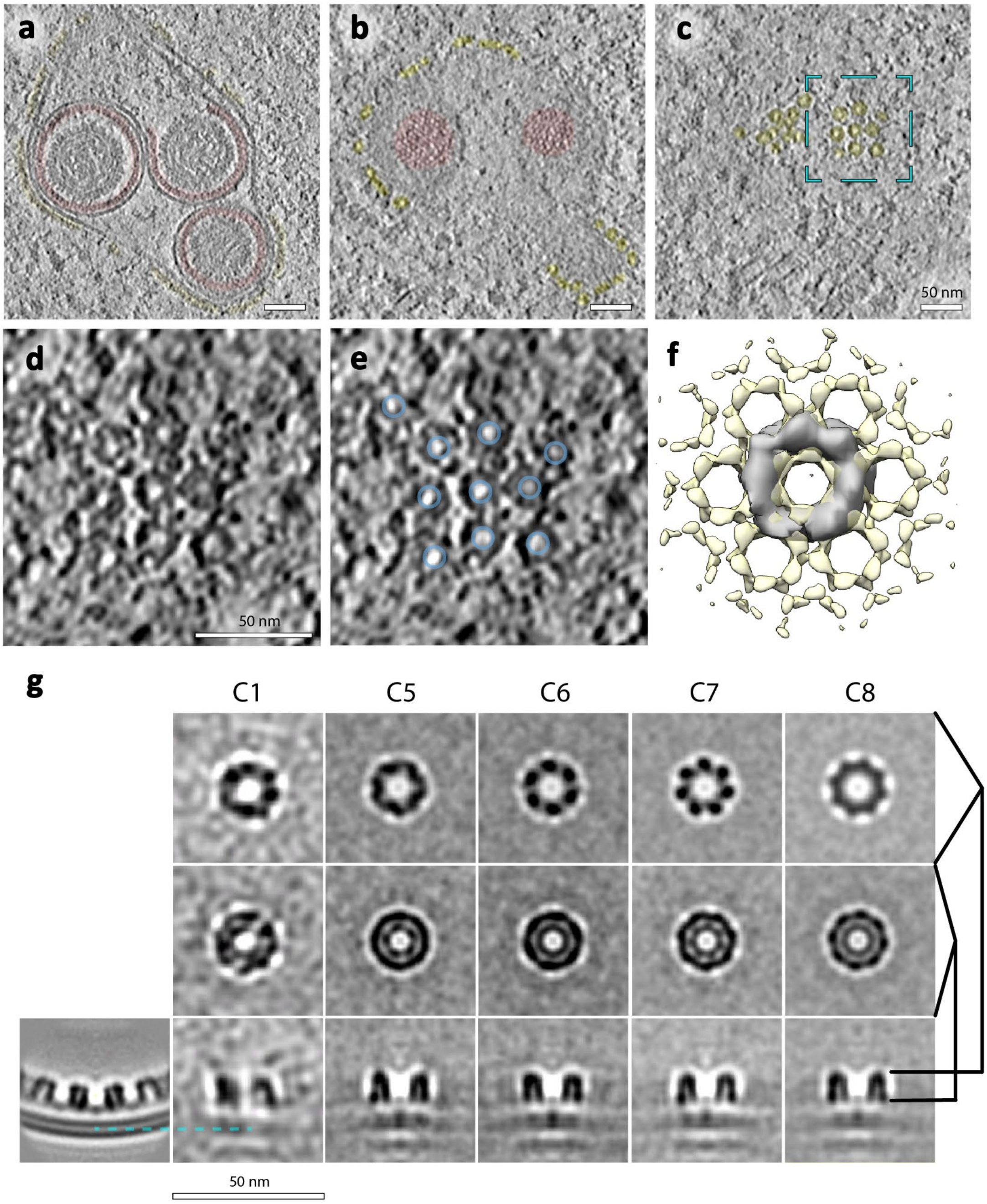


**Figure S8. Putative NEC coat on negatively curved surfaces.** The thickness and distance to the membrane of this layer are consistent with the canonical NEC lattice. **a, b, c** slices through the same nucleoplasmic reticulum at different depths showing the distribution of the putative NEC layer. **d, e** An enlarged section of panel c, showing top views of ring-like structures (highlighted in blue in e). **j** Overlay of the surface representation of the average volume of 121 ring particles from two tomograms and the canonical NEC lattice. Notably, the membrane was not included as an alignment feature. Each ring could plausibly accommodate two concentric layers of pUL31/34 dimers. Averaging a more exhaustive (but less stringently picked) set of negatively curved lattice particles did not converge (and is therefore not shown), suggesting a high degree of variability. **g** Sections through the ring average volume with different C symmetries applied. Visually, C7 is the best match to C1 but it is possible these structures have no strict symmetry, as suggested by d, e. Note that symmetrisation in this case means addition of subvolumes at defined rotations (e.g. 5 subvolumes with 60° degree increments for C6 symmetry). Alignment was performed after the addition of symmetry related particles. The bottom left-most panel is a section through the canonical NEC lattice.

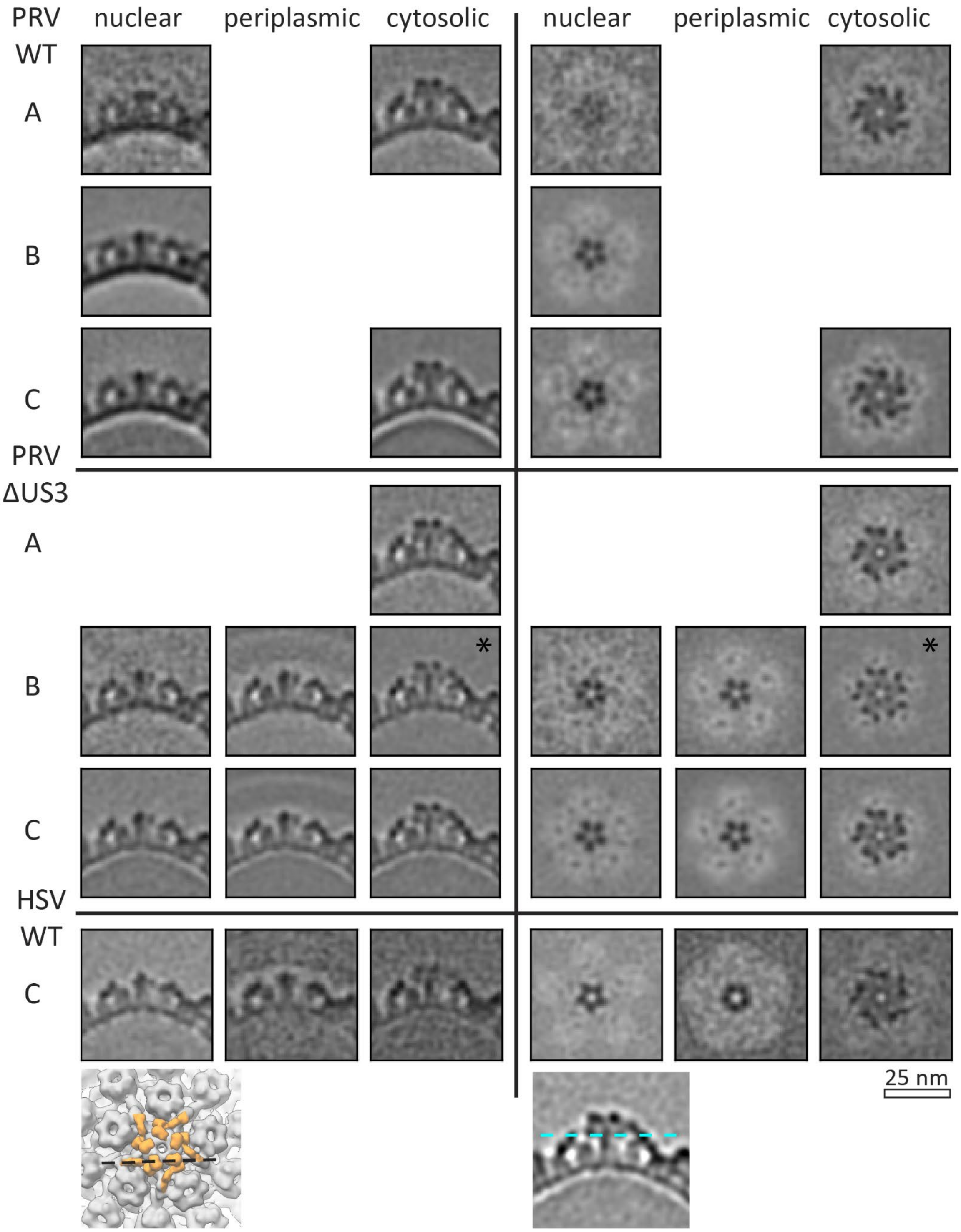


**Figure S9. Overview of capsid volumes.** Shown are normalised volumes of A-, B-, and C- capsids located in the nucleoplasm, perinuclear vesicles, and cytosol. Particles included in the perinuclear B- class consist of capsids with visible scaffolds or partially spooled genomes.

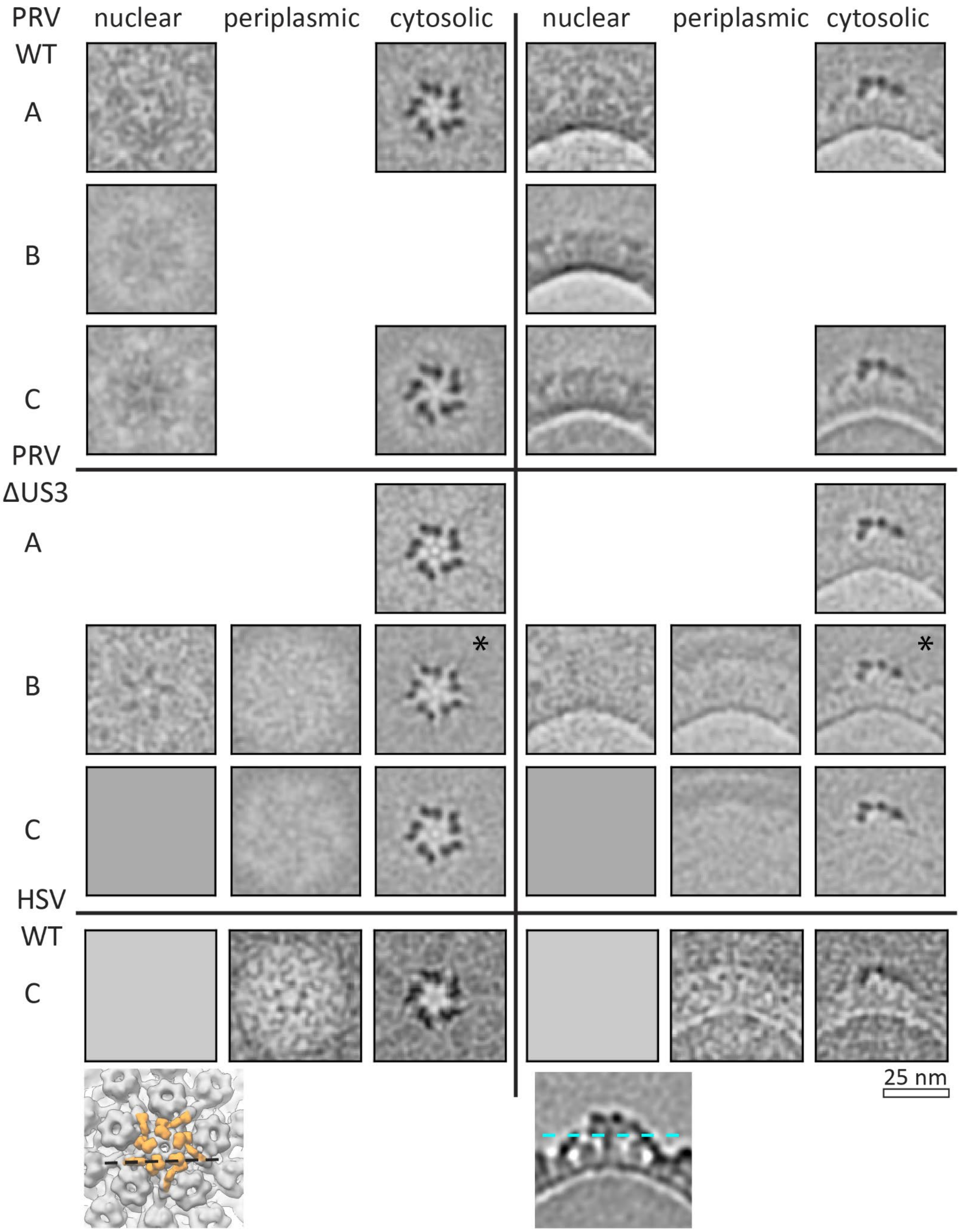


**Figure S10. Overview of normalised capsid volume differences.** PrV-ΔUS3 nuclear C-capsid volume was subtracted from WT PrV and PrV-ΔUS3 A-, B-, and C- capsid volumes derived from particles located in the nucleoplasm, perinuclear vesicles, and cytosol. HSV1 nuclear C-capsid volume was subtracted from HSV perinuclear and nuclear C-capsids.

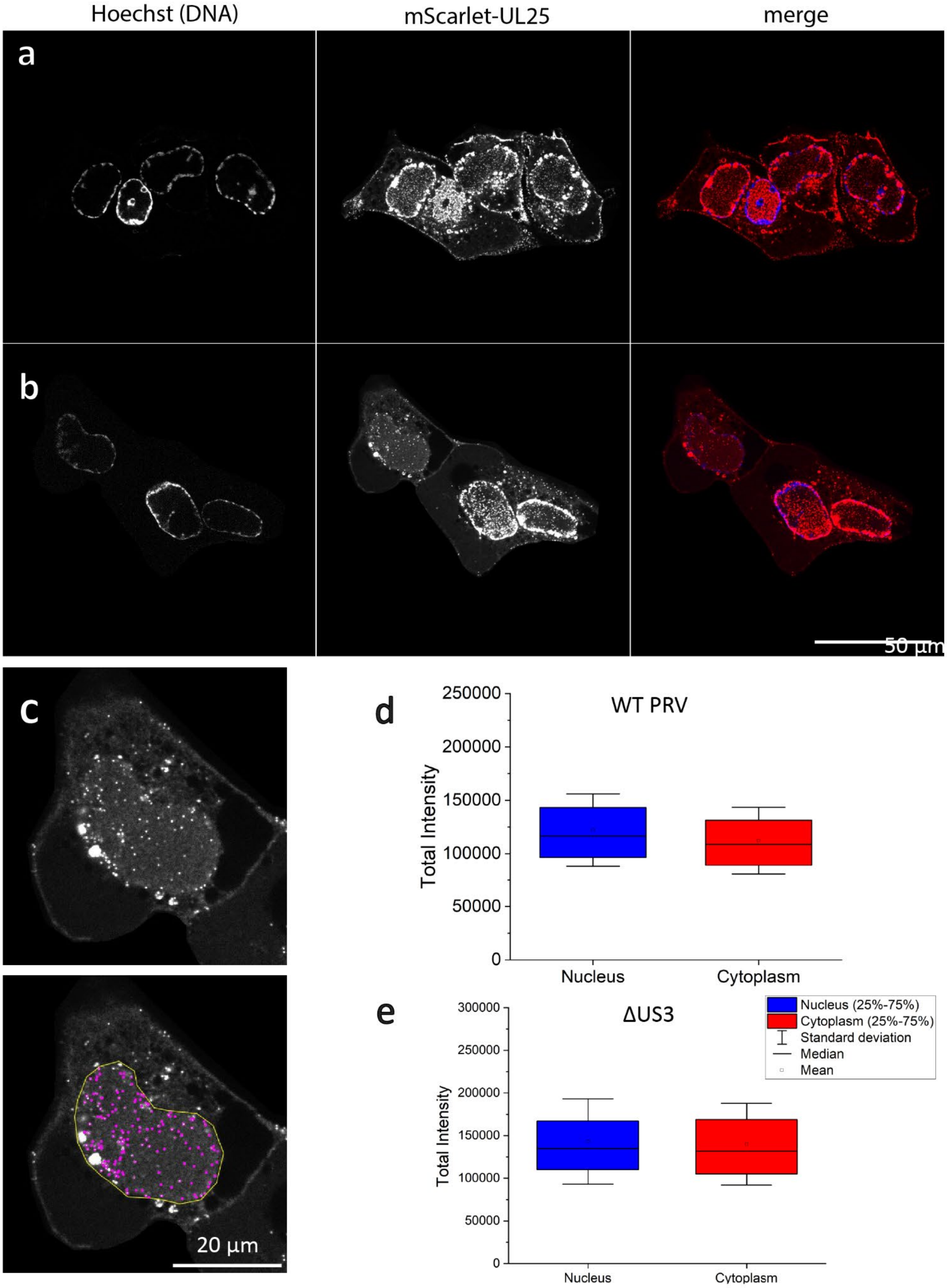


**Figure S11. Localization of individual nuclear and cytoplasmic mScarlet-UL25 labelled capsids. a** PK15 cells were infected with PrV-mScarlet-UL25 or **b** PrV-mScarlet-UL25-ΔUS3 fixed at 7 or 10 hpi, respectively, and imaged using spinning disc microscopy. UL25-mScarlet (red); DNA-Hoechst (blue). One plane of the acquired volume is shown. **c** Viral particles were detected in the 3D volumes using Trackmate in FIJI with an expected blob diameter of 0.4 microns and the quality threshold set to 10.000. Fluorescent signal in a single plane of a PK15 cell infected with PrV-mScarlet-UL25-ΔUS3 and fixed 10 hpi (top). Projection of all detected single particles (purple) of the volume in a nuclear ROI (yellow) onto one plane (bottom). **d, e** The FIJI plugin Trackmate was used to detect and measure individual virus particle fluorescent intensities. For each condition, the total intensity of more than 4.000 particles was quantified and detected in more than 30 different cells.

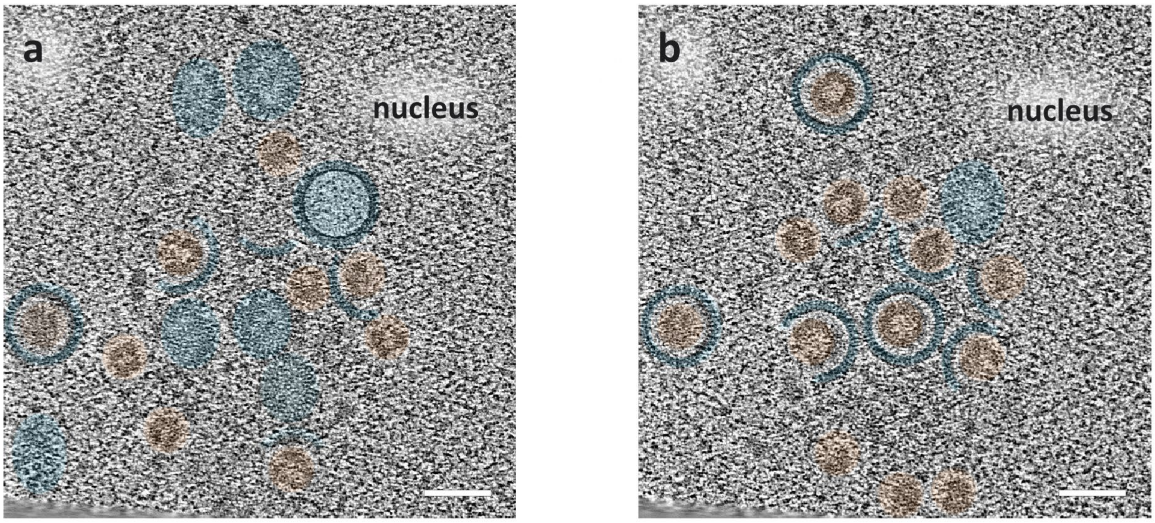


**Figure S12. Nucleoplasm of Vero cells infected with WT PrV at 12 h post infection.** Shown are slices through a tomogram ~25 nm apart in Z. Depicted capsid assembly sites are similar to those in PrV-ΔUS3 (Fig. 1b). Capsid components are highlighted in blue, scaffold rings in orange. Scale bars represent 100 nm.

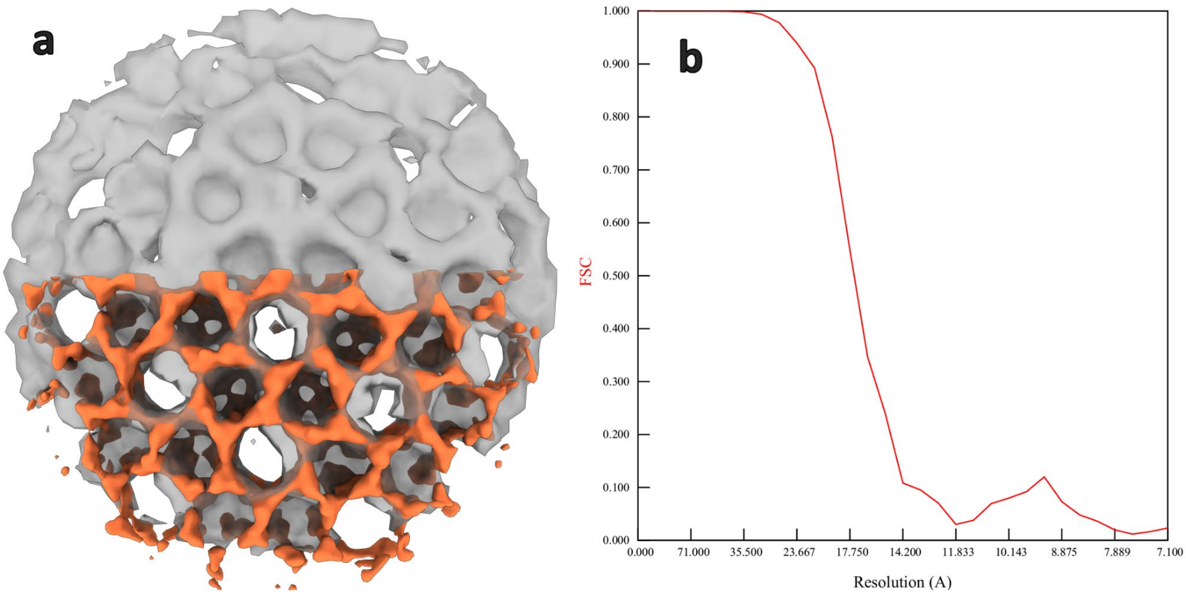


**Figure S13. PrV-ΔUS3 NEC matches the architecture of pUL31/34-GFP. a** Comparison of EMD-3215 (grey) and the map of canonical NEC determined in this study (orange). **b** Gold-standard Fourier-shell correlation of the latter map.

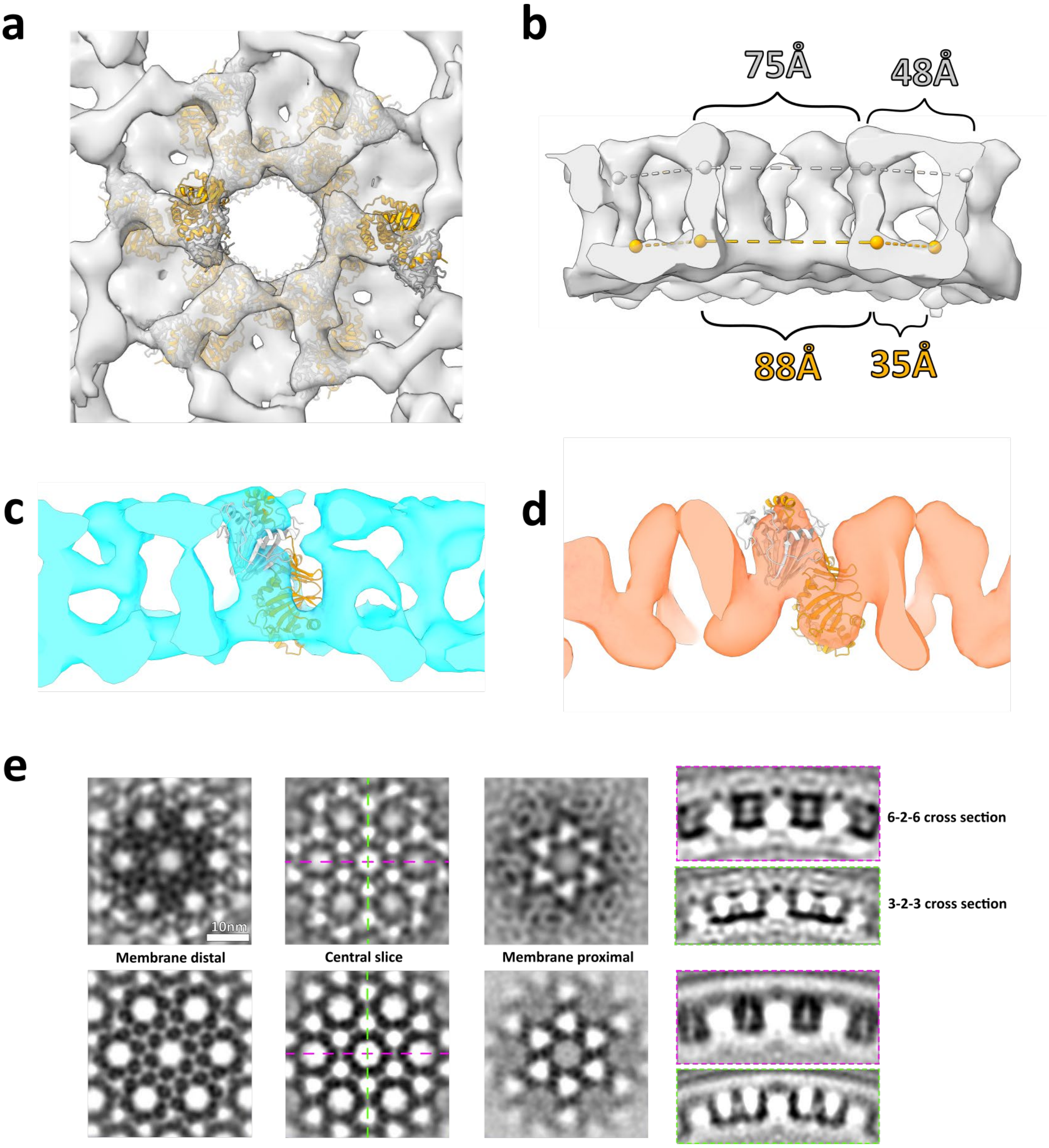


**Figure S14. Fitting of HSV1 pUL31/34 crystal structure into HSV1 NEC map.** The same fitting process was applied as seen in Figure 8 and the Materials and Methods - Rigid Body FItting section. However, the fit led to significant clashes between inter-hexamer dimers, and is therefore even less reliable than the similar PrV analysis. **a** View of the lattice with membrane proximal region facing the viewer. Subvolume average is shown as grey isosurface, pUL31 as orange ribbon and pUL34 as light grey ribbon. **b** Distances between centre of mass markers on opposing sides of the HSV-1 NEC hexamers. Light grey markers represent centre of mass of pUL34 and arm of pUL31 (residues 56-84), orange markers represent centre of mass of the remainder of pUL31. **c** Fit of single HSV-1 pUL31/34 heterodimer into subvolume average of HSV-1 NEC. **d** Fit of single HSV-1 pUL31/34 heterodimer into subvolume average of PrV NEC. **e** Slices through the electron densities of the HSV-1 NEC (top) and the PrV NEC (bottom) subvolume averaging structures. From left to right: membrane distal slice, representing the pUL34 part of each heterodimer; central slice, with the green dotted line along the 6-2-6 direction and the magenta line through the 3-2-3 direction of the lattice; membrane proximal slice, representing pUL34 densities; and finally cross sections through the 6-2-6 and 3-2-3 directions of the NEC.

| PrV WT | | nuclear | perinuclear | cytosolic |
|---|---|---|---|---|
| A | N | 410 | N/A | 325 |
| | FSC 0.143 | 40 | - | 27 |
| | EMD | - | - | - |
| B | N | 5445 | N/A | N/A |
| | FSC 0.143 | 22 | - | - |
| | EMD | - | - | - |
| C | N | 1920 | N/A | 3445 |
| | FSC 0.143 | 29 | - | 26 |
| | EMD | | - | |
| PrV-ΔUS3 | | nuclear | perinuclear | cytosolic |
| A | N | N/A | N/A | 410 |
| | FSC 0.143 | - | - | 34 |
| | EMD | - | - | - |
| B | N | 295 | 1030 | 735 |
| | FSC 0.143 | 31 | 29* | 22 |
| | EMD | - | - | - |
| C | N | 1665 | 3950 | 1065 |
| | FSC 0.143 | 24 | 23 | 31 |
| | EMD | | | |
| HSV WT | | nuclear | perinuclear | cytosolic |
| C | N | 1170 | 205* | 450 |
| | FSC 0.143 | 30 | 38 | 35 |
| | EMD | | | |

Supplemental table 1. Nucleocapsid maps presented in this study. *-mixed type of capsids.

| NEC | | |
|---|---|---|
| Canonical PrV ΔUS3 | N particles | 135,576 |
| | FSC 0.143 | 14 |
| | EMD | |
| Canonical | N particles | 8526 |

| | | |
|---|---|---|
| HSV WT | | |
| | FSC 0.143 | 21 |
| | EMD | |
| Helical<br>PrV ΔUS3 | N particles | 4247 |
| | FSC 0.143 | 21 |
| | EMD | |

Supplemental table 2. NEC maps presented in this study.

### Supplemental movie legends

**Supp. Movie 1 – Capsid assembly, DNA spooling and perinuclear vesicle formation observed in the nucleus.** Tomographic volume of capsids, perinuclear vesicles and other structures observable in the nucleus. Orange arrows heads indicate scaffolds found within chromatin deficient areas. Magenta arrowhead shows a partial capsid that may be undergoing assembly. Yellow arrowhead points towards assembling B-capsids which are yet to spool DNA. Cyan arrowhead shows a capsid in the process of spooling DNA, which is assumed due to the more loosely packed density within the capsid – individual DNA fibres are visible. Blue arrowhead show C-capsids in the nucleus containing densely packed DNA.

**Supp. Movie 2 – Perinuclear vesicle formation and the observation of non-canonical NEC-associated structures.** Tomographic volume of perinuclear vesicles in type-1 nucleoplasmic reticula, capsids budding into perinuclear vesicles and tubular NEC-associated structure. Red arrows point toward assembling heterodimeric NEC arrays at the inner nuclear membrane that have associated with capsids and a vesicle-like structure (pink arrowhead). A green asterisk denotes a perinuclear vesicle that has formed and enveloped 2 individual capsids (an A-capsid and a C-capsid). The blue asterisk denotes an NEC-associated tube, the sub-volume average for which can be seen in Fig. 7. A white arrowhead shows a small opening toward the nucleus in a perinuclear vesicle that is almost fully formed.

**Supp. Movie 3 – Perinuclear vesicles opening to the cytosol of a cell.** Tomographic volume of perinuclear vesicles in the lumen between the inner and outer nuclear membranes. The black arrowheads point towards openings of perinuclear vesicles at the outer nuclear membrane towards the cytosol.

**Supp. Movie 4 – Multiple NEC-associated structures can assemble into Type-1 NR herniations.** Tomographic volume of NEC-associated structures in type-1 nucleoplasmic reticula herniations and invagination of the INM by the NEC. Orange arrows heads represent scaffolds found, once again, within chromatin deficient areas. In one area of the tomogram a scaffold is visible in an NEC tube. Yellow arrowhead points towards assembling B-capsids which are yet to spool DNA while blue arrowheads point toward fully formed C-capsids. Red arrowhead points to invagination sites caused by the NEC. On one of these occasions no structures are apparent to cause such an invagination to occur (bottom). Green asterisk shows a perinuclear vesicle containing B-scaffold. The blue arrow shows an NEC tube-like structure.
